# Anti-Windup Protection Circuits for Biomolecular Integral Controllers

**DOI:** 10.1101/2023.10.06.561168

**Authors:** Maurice Filo, Ankit Gupta, Mustafa Khammash

## Abstract

Robust Perfect Adaptation (RPA) is a desired property of biological systems wherein a system’s output perfectly adapts to a steady state, irrespective of a broad class of perturbations. Achieving RPA typically requires the deployment of integral controllers, which continually adjust the system’s output based on the cumulative error over time. However, the action of these integral controllers can lead to a phenomenon known as “windup”. Windup occurs when an actuator in the system is unable to respond to the controller’s commands, often due to physical constraints, causing the integral error to accumulate significantly. In biomolecular control systems, this phenomenon is especially pronounced due to the positivity of molecular concentrations, inevitable promoter saturation and resource limitations. To protect against such performance deterioration or even instability, we present three biomolecular anti-windup topologies. The underlying architectures of these topologies are then linked to classical control-theoretic anti-windup strategies. This link is made possible due the development of a general model reduction result for chemical reaction networks with fast sequestration reactions that is valid in both the deterministic and stochastic settings. The topologies are realized as chemical reaction networks for which genetic designs, harnessing the flexibility of inteins, are proposed. To validate the efficacy of our designs in mitigating windup effects, we perform simulations across a range of biological systems, including a complex model of Type I diabetic patients and advanced biomolecular proportional-integral-derivative (PID) controllers. This work lays a foundation for developing robust and reliable biomolecular control systems, providing necessary safety and protection against windup-induced instability.

## I. INTRODUCTION

Control theory has long been fundamental to the advancement of engineering systems, providing principles and methods to guide system behavior towards desired outcomes. More recently, these principles have been adopted within the growing field of synthetic biology, leading to the creation of sophisticated, biomolecular feedback controllers capable of executing complex tasks [1]–[4]. In particular, integral feedback controllers (IFCs) play a unique role among these controllers due to their ability to enable robust perfect adaptation (RPA) [5] in the system. RPA is similar to homeostasis, a recurrent principle in biology, but it is even more stringent. In essence, RPA means that a specific variable in the system is always regulated to maintain a fixed steady-state value, referred to as the setpoint, regardless of any constant disturbances or uncertainties that might be present. To this end, the antithetic integral feedback (AIF) controller was introduced to provide a chemical reaction network (CRN) that mathematically realizes integral feedback control via a sequence of chemical reactions [6]. Remarkably, this biomolecular controller proved capable of delivering RPA not only in deterministic settings, but also amid the noise inherent in stochastic settings at the population level. Furthermore, it was also shown that the AIF controller indeed represents the minimal design which is both necessary and sufficient for achieving RPA in noisy environments [7], [8]. Building upon the theoretical groundwork, several genetic implementations of integral and quasi-integral feedback controllers were realized in various settings including *in vivo* [7], [9]– 11], *in vitro* [12], and optogenetically *in silico* [13].

Following the introduction of the AIF controller, several studies were undertaken to elucidate its dynamic performance, stability, inherent trade-offs and tuning [14]– 18]. In order to circumvent the limitations of the AIF controller, a new generation of more advanced controllers was developed. These controllers augment the AIF motif with supplementary circuitry that realizes proportional-integral-derivative (PID) controllers [19]– 23]. This advanced class of controllers has been demonstrated to enhance dynamic performance and reduce noise, embodied as cell-to-cell variability, all while preserving the RPA property.

Despite their promise, there are still challenges that limit the full potential of biomolecular IFCs. Among these challenges, one of the most noteworthy is integral windup [24] – a phenomenon where the integral term in the controller accumulates an error over time, causing the controller to overshoot (or undershoot) the setpoint when the actuation (or sensing) is saturating. This problem can lead to extended periods of poor performance and even system instability. It becomes especially relevant in the context of biomolecular systems where saturation is often encountered due to positivity of molecular concentrations, promoter saturation and limited resources such as enzyme concentrations and energy molecules [25]– [28]. Integral windup is not a challenge exclusive to biomolecular systems. Indeed, its origins and subsequent mitigation strategies can be traced back to the 1930s [29] in industrial applications. Actuators, with their saturating upper and lower operational limits, have long been recognized as major contributors to integral windup manifesting as poor dynamic performance or even instability. In particular, a positivity constraint on actuators is nothing but a special case of actuator saturation where the lower bound is inherently zero. This specific form of saturation, associated with the zero-bound, has been acknowledged for decades as a potential catalyst for instability arising from integral windup. Classic scenarios include valves incapable of being more than fully open or fully closed [30]–[32], or heaters incapable of cooling [33]. Clearly, this phenomenon is universal, reinforcing the pervasive relevance and importance of the integral windup issue across diverse domains, including biology. To this end, when used in practical applications, integral controllers are usually accompanied by anti-windup mechanisms [29].

For a better understanding of integral windup within the framework of biomolecular controllers, consider the visual illustration depicted in Fig. 1(a). This schematic portrays a closed-loop network consisting of an arbitrary process to be regulated and a feedback controller network incorporating an IFC. In this setting, the output variable of interest is species **Y**, which is sensed by the controller network. Subsequently, the controller network operates on the process, aiming to robustly navigate the levels of **Y** towards a specific target setpoint despite constant disturbances and uncertainties in the process. In some cases, the designated setpoint may demand a degree of actuation that exceeds the capacity of the actuator, due to unavoidable saturation constraints. This leads to a situation where the controller persists in attempting to achieve an unattainable level of actuation and, as a result, certain controller species accumulates. In mathematical terms, consider the IFC represented by a variable denoted by 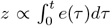, where *e*(*t*) ≜ *r* − *y*(*t*) is the error signal denoting the current deviation of the output from the desired setpoint *r*. If the control action *u*(*t*) reaches saturation, thereby inhibiting the controller’s ability to drive the error to zero, the integral continues to grow, and thus leading to windup. Note that a similar scenario may also happen when the sensor saturates. Therefore, it becomes essential to design a biomolecular protection circuit implementing an anti-windup strategy. This circuit should be capable of intervening to mitigate such phenomena when they occur, while remaining passive when the system is operating within a safe regime. In other words, the controller should be capable of operating in two modes: a normal mode where integral feedback facilitates Robust Perfect Adaptation (RPA) and a protection mode where aiming to achieve RPA is forfeited as a necessary compromise to uphold safety measures.

**Fig. 1:**
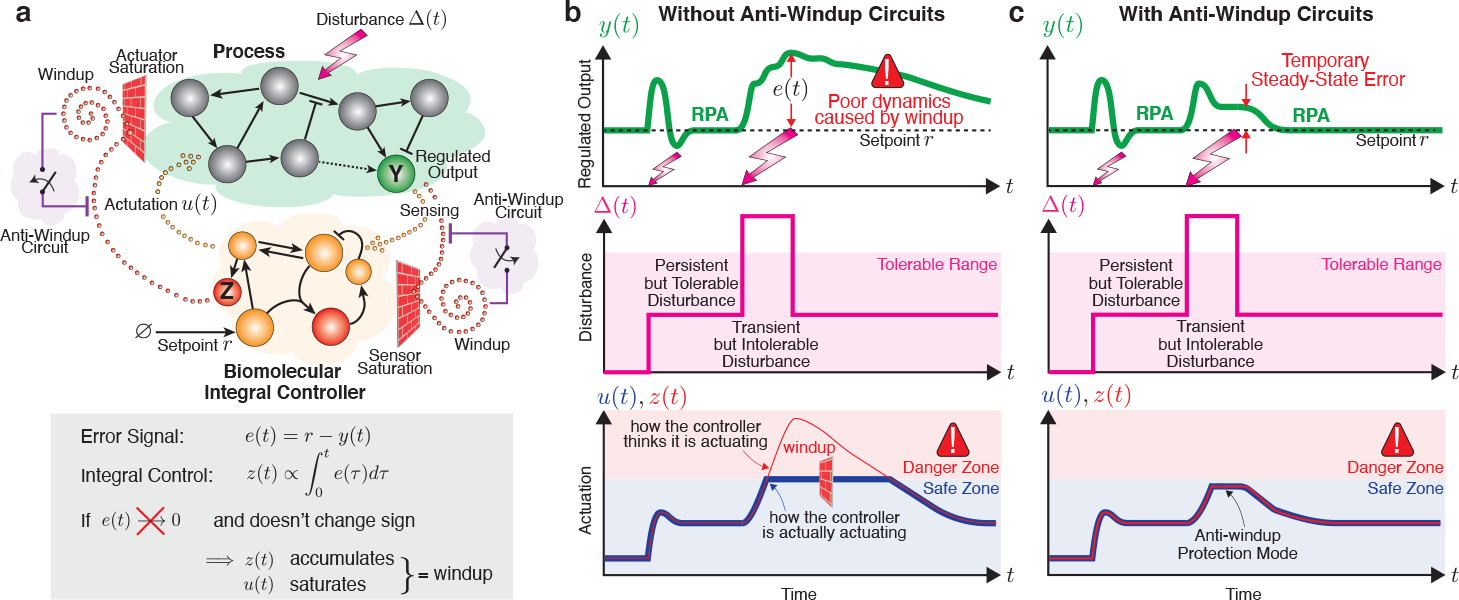
Integral windup. (a) A graphical portrayal of integral windup in a biomolecular system. The closed-loop network is comprised of an arbitrary process in a feedback configuration with a controller network involving integral feedback control. The controller’s objective is to drive the regulated output of interest **Y** to a prescribed setpoint *r* at steady state, despite the presence of constant disturbances and uncertainties – a property referred to as Robust Perfect Adaptation (RPA). The controller essentially operates by sensing the output **Y**, computing the integral of the error *e*(*t*) ≜ *r* − *y*(*t*) and feeding the integral control action back into the process. At steady-state, if it exists, *ż* = 0 and thus the error converges to zero. However, saturations in the actuator and/or sensor might obstruct this process, thereby preventing the error from reducing to zero. This results in the integral of the error accumulating, leading to windup. Such a condition causes certain molecular species, denoted here as **Z**, to increase uncontrollably, as symbolized by the spiral “winding up” due to bouncing off of a saturation wall. Therefore, equipping the controller with anti-windup circuitry becomes indispensable as a protection measure preventing the uncontrolled growth of **Z**. (b) A typical behavior in the absence of anti-windup circuits. Tolerable disturbances ∆ can be promptly rejected by the controller, thus ensuring Robust Perfect Adaptation (RPA). However, when the disturbance exceeds the manageable threshold, actuation *u* hits its saturation limit and a control species **Z** starts to accumulate, triggering windup. Even when the disturbance subsides, the error takes an extended period to return to zero because *z*(*t*) is significantly beyond the safe zone, resulting in a prolonged “unwinding” process. (c) Anti-windup circuits offer a solution to this issue by stepping in only when necessary, thus preventing the controller from reaching a dangerous state. This intervention comes at the cost of a steady-state error which is anyway inevitable. However, it helps in avoiding the windup process and ensuring that once the severe disturbance subsides, the error can quickly return to zero.

Fig. 1(b) graphically demonstrates how windup could occur due to an intense transient disturbance beyond the system’s tolerance. This can lead to a saturation in the control action *u*, resulting in an accumulation in the control variable *z*. Consequently, it causes a persistent error *e* that lingers long after the disturbance has subsided, thus taking an extended period to return to zero. Fig. 1(c) depicts the impact of implementing an anti-windup strategy. Essentially, this strategy prevents the accumulation of the control variable *z* beyond a defined safe threshold by “forgetting” to integrate when operating in the danger zone. As a result, it incurs a non-zero steady-state error but expedites the return to zero error once the disturbance recedes. In a sense, windup could be conceptualized as a “trauma” inflicted on the system due to a severe disturbance. The ramifications of this trauma tend to linger, taking a significant amount of time to fade even after the cause of the trauma has subsided. In this context, an anti-windup strategy can be seen as a therapeutic intervention, helping the controller to move past its “traumatic” experience by promoting a form of forgetfulness towards the disruptive event.

Various anti-windup strategies have been proposed in the literature of traditional control systems [29], [34], [35]. These strategies modify the integrator’s behavior within the controller upon detecting saturation, effectively preventing excessive error accumulation and its detrimental consequences. Nonetheless, implementing these strategies in the context of biomolecular systems poses unique challenges, primarily due to the inherent complexity and constrained structure of nonlinear CRNs. In this paper, we address this challenge by proposing CRN-based biomolecular realization of three anti-windup topologies. We start by phenomenologically modeling the dynamics of each topology to motivate their effectiveness in preventing windup. We then present realizations of these topologies as CRNs. Finally, we propose genetic realizations of the anti-windup circuitry – a step forward towards their implementation in living cells.

## II. Notation

Uppercase bold letters, e.g. **Y**, are reserved for species names. Their corresponding lowercase letters, e.g. *y*(*t*), represent their deterministic concentrations as time-varying signals, where *t* is time. Over-bars, e.g. 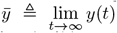 represent steady-state values when they exist. A tilde, e.g. 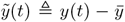, represents the deviation from the steadystate value, and a hat, e.g. *ŷ*(*s*), represents the Laplace transform of 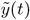, where *s* is the Laplace variable. The *s* and *t* variables are suppressed when they are clear from context. ℝ_+_ and ℝ_−_ denote the set of non-negative and non-positive real numbers, respectively. *e*_*i*_ is a vector, of the appropriate size, whose entries are all zeros except the *i*^th^-entry being 1. Let *f* be a function, then range(*f*) denotes its range. Let of the two functions, that is (*f ° g*)(*x*) = *f g*(*x*). If *z* is *g* be another function, then *f ° g* denotes the composition a scalar or a time varying signal, then *z*^+^ ≜ max(*z*, 0) and *z*^−^ ≜ max(−*z*, 0). Finally, ||.|| denotes the *ℓ*^2^-norm.

## III. Results

We begin by laying out the theoretical foundation, including the introduction of key definitions pertaining to the process to be controlled. We also provide concrete examples to clarify the definitions. We subsequently close the loop using the classical IFC suffering from possible actuator and sensor saturations, and extend this discussion to the biomolecular AIF controller, drawing a rigorous mathematical link to the classical IFC. We then demonstrate how windup may arise with AIF control even in the absence of molecular saturations due to the positive nature of the AIF motif. We show how this observation fits the classical IFC framework. Further, we delve into the occurrence of windup due to molecular saturations and present three different topologies, complemented by suitable CRN realizations, that effectively implement anti-windup strategies. Potential genetic implementations are also proposed. Lastly, we demonstrate the efficacy of our anti-windup designs using three numerical simulations, including an FDA-approved model for glucose-insulin response in diabetic patients (Type I) [36], [37].

### A. Open-Loop System

Consider a general process 𝒫_∆_ to be controlled, depicted in Fig. 2(a), which is modeled as a nonlinear dynamical system with input and output signals *u* and *y*, respectively. The process is parameterized by ∆ which can be thought of as an external disturbance or uncertainty. For simplicity we consider processes that are single-input single-output (SISO) throughout the paper, that is, *u, y*∈ ℝ; however, this can be easily generalized to multiple input multiple output (MIMO) systems. An example of such process 𝒫_∆_ is given by the following nonlinear state-space model

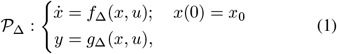

where *x*∈ ℝ^*n*^ represents the internal state vector and *x*_0_ represents the initial condition. Without loss of generality, we will proceed under the assumption that initial conditions are zero, unless specified differently. Let 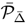 denote the steady-state input/output map of the process. For the nonlinear state-space realization given in (1), the steady-state input/output map is given implicitly as the following set of nonlinear algebraic equations

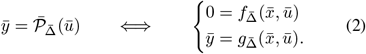

The objective here is to design a feedback controller that endows the output *y* with RPA, that is, it steers *y* to a prescribed steady-state value or setpoint 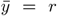, despite the presence of constant disturbances or uncertainties and regardless of the initial conditions.

Let us define four important concepts that are solely related to the controlled process and disturbance represented as 𝒫_∆_. In the following definitions, we denote by 𝕌 and 𝔻 as the set of feasible inputs and disturbances, respectively.

**Fig. 2:**
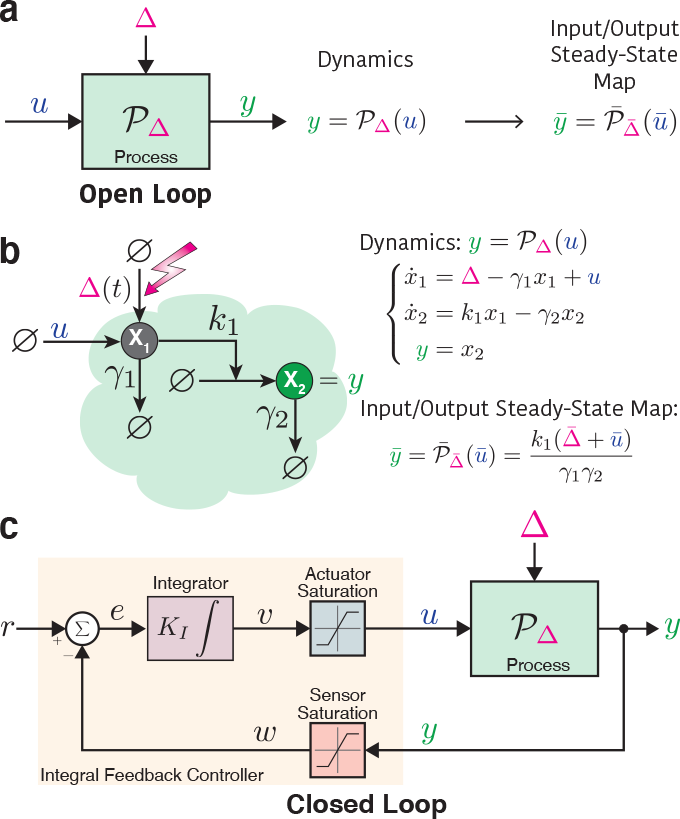
Classical integral feedback control. (a) Process or open-loop system. The input and output of the process are respectively denoted by *u* and *y*, while ∆ denotes a disturbance or uncertainty. The dynamics of the process are described by a (nonlinear) dynamic operator 𝒫_∆_, and its input/output steady-state map for a constant disturbance or uncertainty is denoted by _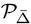_. (b) Gene expression as an example of a simple process. The process is comprised of two molecular species **X**_**1**_ and **X**_**2**_ representing the mRNA and proteins whose degradation rates are denoted by *γ*_1_ and *γ*_2_, respectively. Note that the translation rate is denoted by *k*_1_ and ∆ represents a disturbance entering the dynamics as a basal transcription rate. The input to this open-loop system is an induced transcription rate denoted by *u*, while the output is the protein concentration *y* = *x*_2_. The dynamics and input/output steady-state map are given here. (c) Closing the loop with the classical IFC. Ideally, the integral controller senses the output *y* and computes the integral of the error *e*(*t*) ≜ *r* − *y*, which signifies the deviation of the output from the setpoint *r*, multiplies this integral by a gain *K*_*I*_, and feeds the result back to the process through the actuator. In more practical scenarios, however, sensor and actuator saturation may occur, impeding the faithful transmission of signals between the controller and the process. The equations describing the dynamics of the closed-loop system are presented in (9).

#### Definition 1 (Supporting Input)

The supporting input, if it exists, for a given desired setpoint *r* and a disturbance 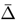 is the input value that will result in a steady-state output value equal to the setpoint *r*. That is, the supporting input *ū* satisfies 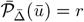, and thus 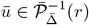.

#### Definition 2 (Admissible Setpoint)

A given setpoint *r* is admissible if it admits a feasible supporting input. That is, for a given setpoint *r* and disturbance or uncertainty 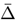, ∃ a feasible supporting input *ū* ∈ 𝕌 such that 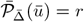.

#### Definition 3 (Set of Admissible Setpoints)

For a given process 𝒫 _∆_ and a set of feasible inputs 𝕌, the set of admissible setpoints, denoted by ℛ (𝒫 _∆_, 𝕌), is the set of all possible admissible setpoints. That is,

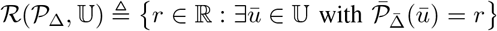

#### Definition 4 (Set of Admissible Disturbances)

For a given admissible setpoint *r*, the set of admissible disturbances, denoted by 𝒟 _*r*_(𝒫 _∆_, 𝕌, 𝔻), is the range of all feasible disturbances that do not destroy the admissibility of the desired setpoint *r*. That is,

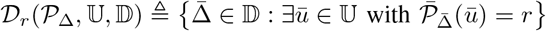

Note that we drop the arguments of ℛ and 𝔻 _*r*_ whenever they are clear from context. Before we provide examples, we make three remarks on the introduced definitions.

#### Remark 1

Admissibility is a concept that depends solely on the process to be controlled, the actuation mechanism (e.g. production, removal, saturation) and disturbance or uncertainty; it is completely independent of the controller structure. Hence, if for a given actuation mechanism and disturbance, a desired setpoint *r* ∉ *ℛ* is not admissible, it means that there is absolutely no controller (regardless of its structure) that can deliver such a setpoint.

#### Remark 2

Admissibility is an algebraic steady-state property since it is ultimately related to the solvability of a set of algebraic equations. Hence, admissibility is a concept that is independent of stability and, as such, if a setpoint is admissible, the dynamics might still be unstable.

#### Remark 3

Throughout the paper, we have, for the sake of simplicity, presumed that for any given desired setpoint *r*, the corresponding supporting input 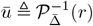, if it exists, is unique. This assumption can be readily relaxed because the exact value of the supporting input is not of primary importance, as long as it is admissible and results in a stable closed-loop fixed point with an output coordinate 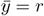.

#### Example 1 (Gene Expression)

Consider the simple model for gene expression given in Fig. 2(b). The actuation and disturbance here take the form of production rates *u* and ∆, respectively. Both of which are assumed to have no saturation. Consequently, only non-negative inputs and disturbances are feasible, that is 𝕌 = 𝔻 = ℝ_+_. The steady-state input/output map is affine in the input *ū* as depicted in Fig. 2(b). As a result, for a given desired setpoint 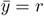, the supporting input is 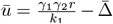. Requiring the supporting input to be feasible (*ū* ≥ 0) yields a condition on the setpoint given by 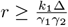. Consequently, the set of admissible setpoints is given by

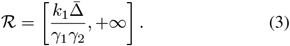

Clearly if there is no basal transcription 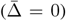, then all non-negative setpoints are admissible. Finally, for a given setpoint *r*, the set of admissible disturbances is calculated by searching for the disturbances that preserve feasibility of the supporting input (*ū* ≥ 0) to obtain

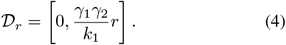

#### Example 2 (Stable Unimolecular Networks)

Consider a general unimolecular network of *L* species, denoted by **X** ≜ {**X**_**1**_, **X**_**2**_, · · ·, **X**_**L**_}, reacting among each other via *K* reaction channels. Let *S*_∆_ and *λ*_∆_(.) respectively denote the stoichiometry matrix and propensity function of the network in the absence of an external actuation, parameterized by a disturbance or uncertainty ∆. For unimolecular reactions, the propensity function is affine in the species concentrations, that is *λ*_∆_(*x*) ≜ *W*_∆_*x* + *b*_∆_, where *W*_∆_ is a *K × L* matrix and *b*_∆_ is a *K ×* 1 vector with non-negative entries. Without loss of generality, let the output of interest be **X**_**L**_, and the actuated input be **X**_**1**_ where the actuation takes the form of a production reaction, that is 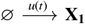, and so 𝕌 = ℝ.

The dynamics are thus given by

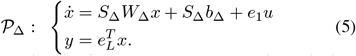

Let us assume that the dynamics are Hurwitz stable, which is equivalent to assuming that the eigenvalues of *S*_∆_*W*_∆_ have strictly negative real parts over a range of disturbances or uncertainties ∆. Then the steady-state input/output map 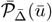 is calculated by setting the time derivative in (5) to zero to obtain

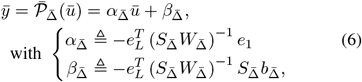

where 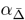is the “steady-state gain” of the network in the absence of basal expressions 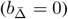, and 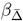 is the “basal offset” of the network that reflects the propagation of the basal expression rates 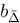. Note that 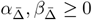. This arises from the fact that 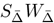 is a Metzler and Hurwitz matrix which means that all entries of its inverse are non-positive [38]. Clearly, the steady-state input/output map is an affine function. As a result, for a given desired setpoint 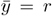, the supporting input is 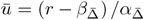. For the setpoint to be admissible, the supporting input needs to be feasible, that is *ū* ∈ 𝕌 = ℝ^+^, and hence the setpoint is required to satisfy 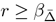. Consequently, the set of admissible setpoints is given by

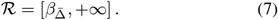

This means that for the setpoint to be admissible it needs to be higher than the basal offset dictated by the non-actuated process. Finally, for a given setpoint *r*, the set of admissible disturbances is

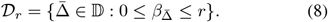

Clearly, Example 1 is a special case where 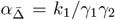 and 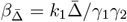.

### B. Closing the Loop with the Classical Integral Controller

According to the internal model principle [39], RPA can be achieved using integral feedback controllers. To this end, consider the closed-loop system depicted in Fig. 2(c), where the controlled process is now in a feedback interconnection with an integral controller. Ideally, the actuator should faithfully transmit the controller’s output to the controlled process (i.e. *u* = *v*), and the sensor should faithfully transmit the output to the summation junction (i.e. *w* = *y*) to compute the deviation (or error) *e* of the *sensed* output *w* from the desired setpoint *r*. However, practically, the actuator and sensor may saturate beyond certain thresholds. Even worse, they may deform the actuation and sensed signals. Sensor and actuator saturations are modeled by the saturation blocks depicted in Fig. 2(c). Finally, integral feedback control is achieved by the integrator module which computes the time integral of the error to yield the feedback signal *v*(*t*) to the actuator.

The full dynamics of the closed-loop system are given by

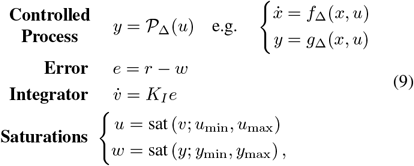

where [*u*_min_, *u*_max_] and [*y*_min_, *y*_max_] represent the actuator and sensor ranges, respectively, and sat(.) is the saturation function defined, for any *z, a, b* ∈ ℝ with *a < b*, as

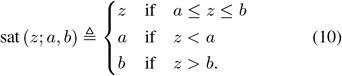

#### Ideal Setting

In the ideal setting, we have the following assumptions:

##### Assumption 1 (Full Admissibility)

All setpoints are admissible by the controlled process for all disturbances or uncertainties, i.e. ℛ= 𝕌 = 𝔻_*r*_ = ℝ, ∀ *r* ∈ ℝ.

##### Assumption 2 (No Saturations)

The sensor and actuator do not exhibit any saturations, i.e. *u*_min_ = *y*_min_ = −∞ and *u*_max_ = *y*_max_ =∞.

Under Assumptions 1 and 2, we have

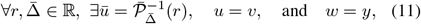

and, as a result, the dynamics given in (9) boils down to

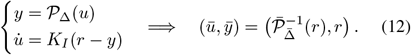

This guarantees that the output *y* will converge at steady-state to the setpoint *r*, independent of 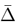, as long as the closed-loop system remains stable, that is RPA is achieved.

#### Non-Ideal Setting

The fixed point of the dynamics of the non-ideal setting can be computed by setting the time derivatives in (9) to zero to obtain the following set of algebraic equations

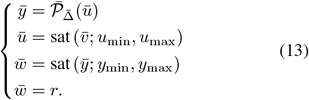

The following lemma characterizes the existence of the fixed point.

##### Lemma 1

Consider the closed-loop system depicted in Fig. 2(c), where the actuator and sensor ranges are given as [*u*_min_, *u*_max_] and [*y*_min_, *y*_max_], respectively. For a given desired setpoint *r* and a steady-state disturbance or uncertainty 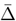, the closed-loop fixed point exists with a feasible supporting input *ū* ∈ 𝕌 if and only if:

- The setpoint is admissible, i.e. *r* ∈ *ℛ*.
- The sensor does not saturate at steady-state, i.e. *r* ∈ [*y*_min_, *y*_max_]
- The actuator does not saturate at steady-state, i.e.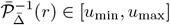.

Furthermore, the fixed point is given by

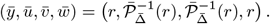

The proof can be found in the Supplementary Information Section S1.

Lemma 1 essentially says that if the desired setpoint is inadmissible by the process, or if either the sensor or actuator saturates, then the integral controller fails and the closed-loop dynamics exhibit no equilibrium. In fact, the only way that integral control fails in achieving RPA is by losing stability of the closed loop whether this instability is due to nonexistence of a fixed point or the existence of an unstable or unreachable fixed point.

### C. A Model Reduction Result for CRNs

In order to use Lemma 1 in a biological setting, we need to extend it into the realm of CRNs. We first present two theorems that investigate the dynamics and fixed-point behaviors of complex CRNs involving fast sequestration reactions. The versatility and depth offered by these theorems will be harnessed throughout this study.

Consider the two dynamical systems, 𝒮_*η*_ and 𝒮, depicted pictorially in Fig. 3 and described by the following ordinary differential equations

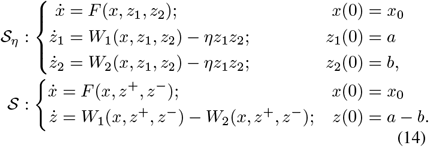

Note that the functional forms of *W*_1_ and *W*_2_ are kept general to include possible degradation terms of *z*_1_ and *z*_2_, among other possible interactions. We make the following assumption.

*Assumption 3:* The three functions *W*_1_, *W*_2_ : 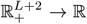 and *F* : 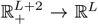 are assumed to be *globally Lipschitz* on their domains, and their form is such that the solution of 𝒮_*η*_ lies in 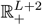.

The following theorem states the convergence of the dynam-ics, as *η* → ∞, of the full system 𝒮_*η*_ to those of the reduced system 𝒮 over any compact time interval.

**Fig. 3:**
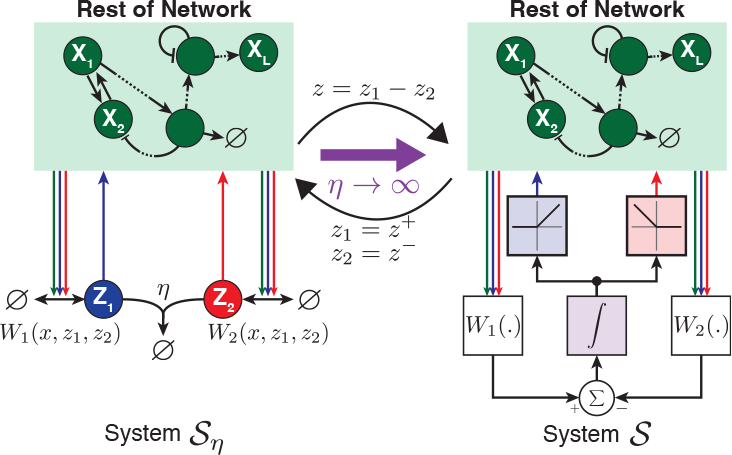
Model reduction for CRNs with fast sequestration reactions. The left schematic illustrates the full system 𝒮_*η*_, where *η* denotes a sequestration rate. In contrast, the right schematic portrays the reduced system 𝒮 which accurately reflects the dynamics of 𝒮_*η*_ as *η* → ∞. Both systems’ precise dynamics are detailed in (14). The system 𝒮_*η*_ consists of two species, **Z**_**1**_ and **Z**_**2**_. Their production and removal rates are compactly represented by the functions *W*_1_ and *W*_2_, respectively. The rest of the network (green box), is arbitrary and remains unchanged in the reduced model. Essentially, when the sequestration rate is fast, the two species operate as a subtraction mechanism, given by *W*_1_ − *W*_2_ which is subsequently integrated in time to produce *Z* = ∫ (*W*_1_ − *W*_2_). Finally, the positive and negative portions of *Z* are isolated and relayed to the rest of the network in place of **Z**_**1**_ and **Z**_**2**_, respectively.

#### Theorem 1

Under Assumption 3 and as *η* → ∞, 𝒮_*η*_ converges to 𝒮 in the following senses: ∀*T >* 0 we have

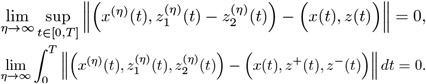

The proof can be found in the Supplementary Information Section S2, and a numerical demonstration is provided in the Supplementary Information, Fig. S4.

#### Remark 4

In the limit as *η* → ∞, the concentration of at least one of the two species **Z**_**1**_ or **Z**_**2**_ is zero at any time. Hence, the dynamics may enter a low-copy regime where stochastic fluctuations become more relevant. To this end, we prove in the Supplementary Information Section S6 that Theorem 1 is also valid in the stochastic setting. We also provide numerical validations in the Supplementary Information, Fig. S5.

The following theorem, on the other hand, states the convergence of the fixed points.

#### Theorem 2

Under Assumption 3 and as *η* → ∞, the limit point 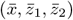 of any convergent sub-sequence of non-negative fixed points of system 𝒮_*η*_ can be transformed to a fixed point of system 𝒮 given by 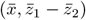.

The following corollary characterizes the relationship between the existence of the fixed points of systems 𝒮_*η*_, as *η* → ∞and 𝒮.

#### Corollary 1

Under Assumption 3, if system 𝒮 admits a unique fixed point 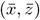, then all convergent sub-sequences of non-negative fixed points of system 𝒮_*η*_ have a single limit point as *η* → ∞ given by 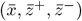. In particular, if there is a bounded set containing all non-negative fixed points of system 𝒮_*η*_, then all the sub-sequences are guaranteed to converge to the unique limiting point 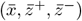.

However, if system admits no fixed points, then there is no convergent sub-sequence of non-negative fixed points of system 𝒮_*η*_. In fact, all sub-sequences will either diverge or converge to a limit fixed point with negative components. In particular, there cannot exist a bounded set containing non-negative fixed points of system 𝒮_*η*_ as *η* → ∞.

The proofs can be found in the Supplementary Information Sections S3 and S4.

### D. AIF Control

Owing to the importance of RPA in biology, the AIF controller was introduced in [6] to realize integral control as a CRN which endows the system with the desirable RPA properties. In this section, we establish a connection between the AIF controller and the classical IFC of Fig. 2(c).

The basic AIF controller is depicted in the closed-loop network of Fig. 4(a), where the objective is to endow the regulated output **X**_**L**_ with RPA. The closed-loop network is constituted of the AIF controller, comprised of two species **Z**_**1**_ and **Z**_**2**_, connected in a feedback configuration with an arbitrary network or process, comprised of *L* species **X**_**1**_, **X**_**2**_, · · ·, **X**_**L**_. For generality, we adopt the biomolecular control paradigm introduced in [20], where we assume that the controller can only interact with the process species **X**_**1**_ and **X**_**L**_ for actuation and sensing, respectively. The controller network involves four reaction channels listed in Table 1.

**TABLE 1:**
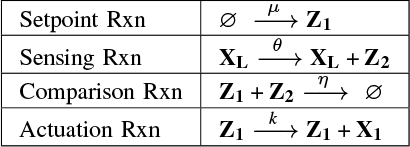
Reactions describing the basic AIF controller.

|  |  |
| --- | --- |
| Setpoint Rxn | $\emptyset \xrightarrow{\mu} \mathbf{Z}_1$ |
| Sensing Rxn | $\mathbf{X}_L \xrightarrow{\theta} \mathbf{X}_L + \mathbf{Z}_2$ |
| Comparison Rxn | $\mathbf{Z}_1 + \mathbf{Z}_2 \xrightarrow{\eta} \emptyset$ |
| Actuation Rxn | $\mathbf{Z}_1 \xrightarrow{k} \mathbf{Z}_1 + \mathbf{X}_1$ |

**Fig. 4:**
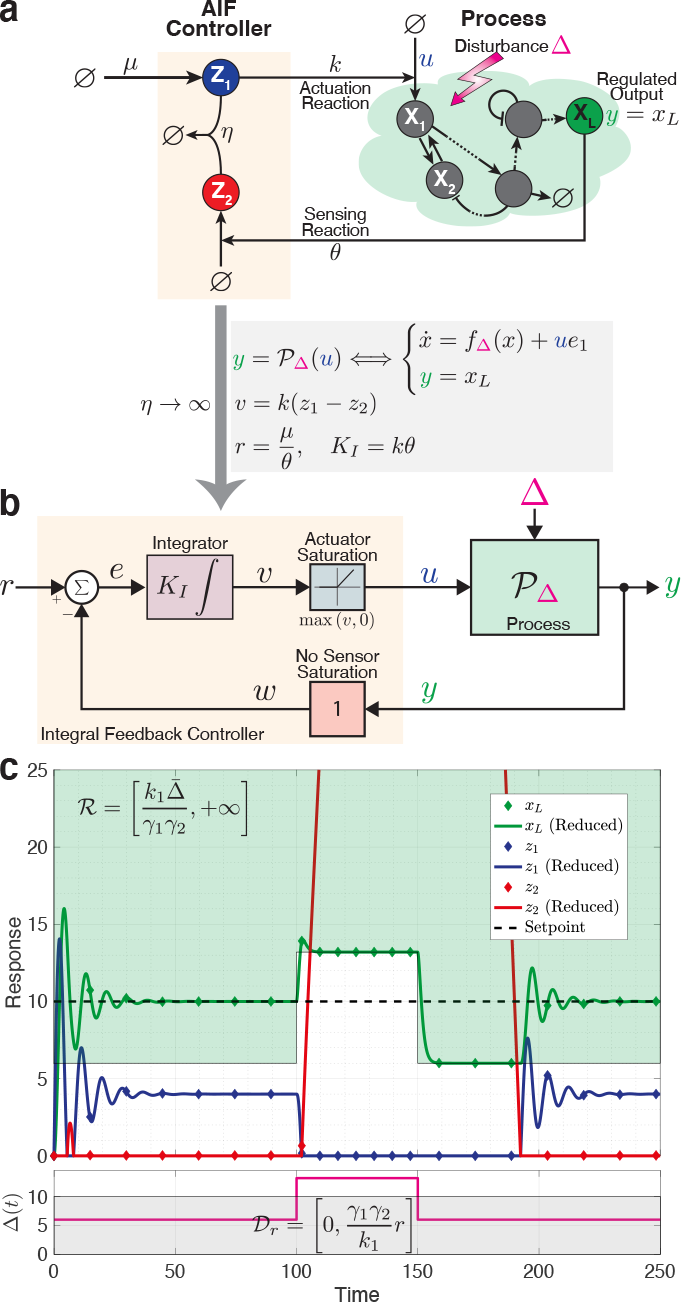
Antithetic integral feedback control. (a) An arbitrary process regulated by the AIF controller. The process is composed of *L* species **X**_**1**_ through **X**_**L**_, with **X**_**L**_ being the regulated output of interest. The AIF controller is composed of two species, **Z**_**1**_ and **Z**_**2**_, which sequester each other at a rate *η*. Sensing is realized via the catalytic production of species **Z**_**2**_ by the output **X**_**L**_ at a rate *θx*_*L*_; whereas, actuation is realized via the catalytic production of the input species **X**_**1**_ at a rate of *u* = *kz*_1_. The setpoint is encoded in the constitutive production of **Z**_**1**_ at a rate *µ*. (b) Block diagram describing the exact closed-loop dynamics in the limit as *η* → ∞. By defining the setpoint *r* and integral gain *K*_*I*_, and introducing an intermediate variable *v*, this diagram can be generated. The diagram demonstrates that, in the strong sequestration limit, the AIF controller is exactly the classical IFC depicted in Fig. 2(c), excluding sensor saturation but including a one-sided actuator saturation due to the inherent positivity of the AIF motif. (c) Integral windup induced by the positivity of the AIF controller. In this simulation example, the process is considered to be the gene expression model presented in Fig. 2(b). The green and gray shaded areas represent the set of admissible setpoints and disturbances _*r*_, respectively (see Example 1 for more details). As the simulation demonstrates, the AIF controller successfully maintains RPA as long as disturbances remain within admissible limits. However, if the disturbance shift to an inadmissible level, the AIF controller falls short, exhibiting poor dynamic performance even after the disturbance returns to an admissible level – a clear demonstration of integral windup. Numerical values: *µ* = 10, *η* = 100, *k* = *θ* = *γ*_1_ = *γ*_2_ = *k*_1_ = 1 and Δ is varied.

The closed-loop dynamics are thus given by

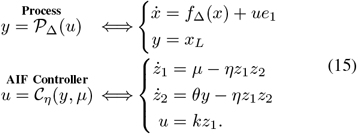

Note that since the actuation by the controller is carried out via a production reaction only which is assumed to have no saturation limit, the set of feasible inputs is 𝕌 = ℝ_+_. The integral action can be seen by looking at the dynamics of *z* ≜ *z*_1_ − *z*_2_ given by

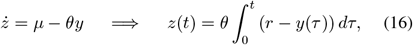

where *r* ≜ *µ/θ* denotes the setpoint. As a result, as long as closed-loop stability is maintained, the regulated output converges to the setpoint 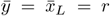 regardless of the initial conditions and process uncertainties/disturbances Δ. As such, the AIF controller endows the regulated output species **X**_**L**_ with RPA.

To reveal the connection between the AIF controller and the classical IFC depicted in Fig. 2(c), we examine the case where the sequestration rate is large enough. In fact, in the limit as *η* → ∞, applying Theorem 1 (with *W*_1_(*x, z*_1_, *z*_2_) ≜ *µ*, and *W*_2_(*x, z*_1_, *z*_2_) ≜ *θx*_*L*_) yields a reduced model for the controller given by

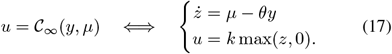

Introducing the intermediate variables *v* ≜ *kz, w* ≜ *y* and *e* ≜ *r* − *w* and the integral gain *K*_*I*_ ≜ *kθ* allows us to draw the block diagram of the closed-loop system depicted in Fig. 4(b). This block diagram allows us to directly compare the AIF controller with the classical IFC depicted in Fig. 2(c). Indeed, the AIF controller is essentially the same as the classical IFC where the sensor has no saturation limits, that is [*y*_min_, *y*_max_] = [−∞, +∞], while the actuator saturates only from below at zero, that is [*u*_min_, *u*_max_] = [0, + ∞]. This is intuitive since the actuation is carried out via the production of **X**_**1**_, and production cannot be negative. As a result, if the desired setpoint *r* requires a supporting input 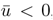, the closed-loop fixed point with the reduced controller 𝒞_∞_ will cease to exist. However, for the full model with large *η*, the closed-loop fixed point may not cease to exist, but instead suffer from a negative component *z*_1_ *<* 0 (see Corollary 1) rendering it unreachable (recall that the positive orthant is invariant under the dynamics of CRNs), and as a result, the dynamics become unstable. One can think of the AIF controller 𝒞_*η*_ with finite *η* as an integral controller coupled with “dynamic saturation” from below at zero, as compared to the case with infinite *η* where the saturation max(*z*, 0) becomes static.

Note that if one has the luxury of changing the actuation mechanism or even adding more actuation mechanisms, the issue of non-existent or unreachable fixed point can be circumvented. For instance, by actuating via both production and degradation of **X**_**1**_, the control action in (15) becomes *u* = *kz*_1_ − *F* (*z*_2_, *x*_*L*_)*x*_1_, and the set of feasible inputs and actuator range become 𝕌 = ℝ. Notably, for *F* (*z*_2_, *x*_*L*_) = *γz*_2_ or *F* (*z*_2_, *x*_*L*_) = *γx*_*L*_, we obtain (filtered) proportional-integral controllers (see [20]).

We close this section, by providing a numerical demonstration depicted in Fig. 4(c) where the gene expression model presented in Fig. 2(b) is controlled by the AIF controller. The plot shows that RPA is achieved as long as the disturbance Δ ∈ 𝔻_*r*_ is admissible. However, when Δ transitions to an inadmissible level, although transiently, integral windup manifests. In this scenario, *z*_1_ attempts to go negative, but of course cannot, due to the positivity of the system. Consequently, *z*_2_ accumulates and remains high for an extended period, even after the disturbance reverts to an admissible level. This results in poor dynamic performance induced by integral windup.

### E. AIF Control with Actuation/Sensing Saturation

Consider now the AIF controller where catalytic production reactions may saturate as depicted in the closed-loop network of Fig. 5(a). The controller network here involves the same four reaction channels described in Table I, but the propensity functions of the actuation and sensing reactions are now replaced with *kh*_*a*_(*z*_1_) and *θh*_*s*_(*x*_*L*_), respectively, where *h*_*a*_ and *h*_*s*_ are nonlinear functions that may introduce saturation. Examples of such functions are Michaelis-Menten or Hill-type functions. The closed-loop dynamics are thus given by

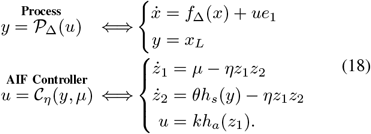

The integral action can still be seen by looking at the dynamics of *z* ≜ *z*_1_ − *z*_2_ given by

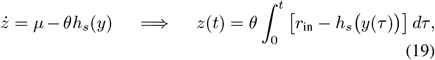

where, *r*_in_ ≜ *µ/θ*. Observe that, assuming closed-loop stability, 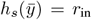. As a result, the sensing function *h*_*s*_ determines the relationship between the input and output setpoints denoted by *r*_in_ and *r*_out_, respectively. Note that in the ideal setting of Fig. 4, *h*_*s*_ is the identity function and thus the input and output setpoints match, that is *r*_in_ = *r*_out_ = *r*. However, in general, the input and output setpoints satisfy the following algebraic equation

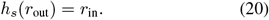

To this end, we refine the concept of admissible setpoints into two distinct categories: admissible input setpoints, denoted as ℛ_in_, and admissible output setpoints, denoted as ℛ_out_.

These can be mapped to each other using Eq. (20) for monotonically increasing functions *h*_*s*_, i.e. ℛ_in_ ≜ *h*_*s*_(_out_). To reveal the connection with the classical IFC depicted in Fig. 2(c), we examine, once again, the case where the sequestration rate is large enough. In fact, in the limit as *η* → ∞, applying Theorem 1 (with *W*_1_(*x, z*_1_, *z*_2_) ≜ *µ* and *W*_2_(*x, z*_1_, *z*_2_) ≜ *θh*_*s*_(*x*_*L*_)), yields a reduced model for the controller given by

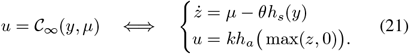

Introducing the intermediate variables *v* ≜ *kz, w* ≜ *h*_*s*_(*y*) and *e* ≜ *r* − *w* and the integral gain *K*_*I*_ ≜ *kθ* allows us to draw the block diagram of the closed-loop system depicted in Fig. 5(b). A direct comparison with the classical IFC, depicted in Fig. 2(c), reveals that the difference here is embodied in the actuator and sensor modules that are now replaced with the nonlinear functions *ψ*_*a*_ and *ψ*_*s*_ define in Fig. 5(a). These functions not only act as saturation components but also deform the signals. The following lemma slightly generalizes Lemma 1 to the case where the saturation blocks of Fig. 2(c) are replaced with the monotonically increasing functions *ψ*_*a*_ and *ψ*_*s*_ shown in Fig. 5(b).

**Fig. 5:**
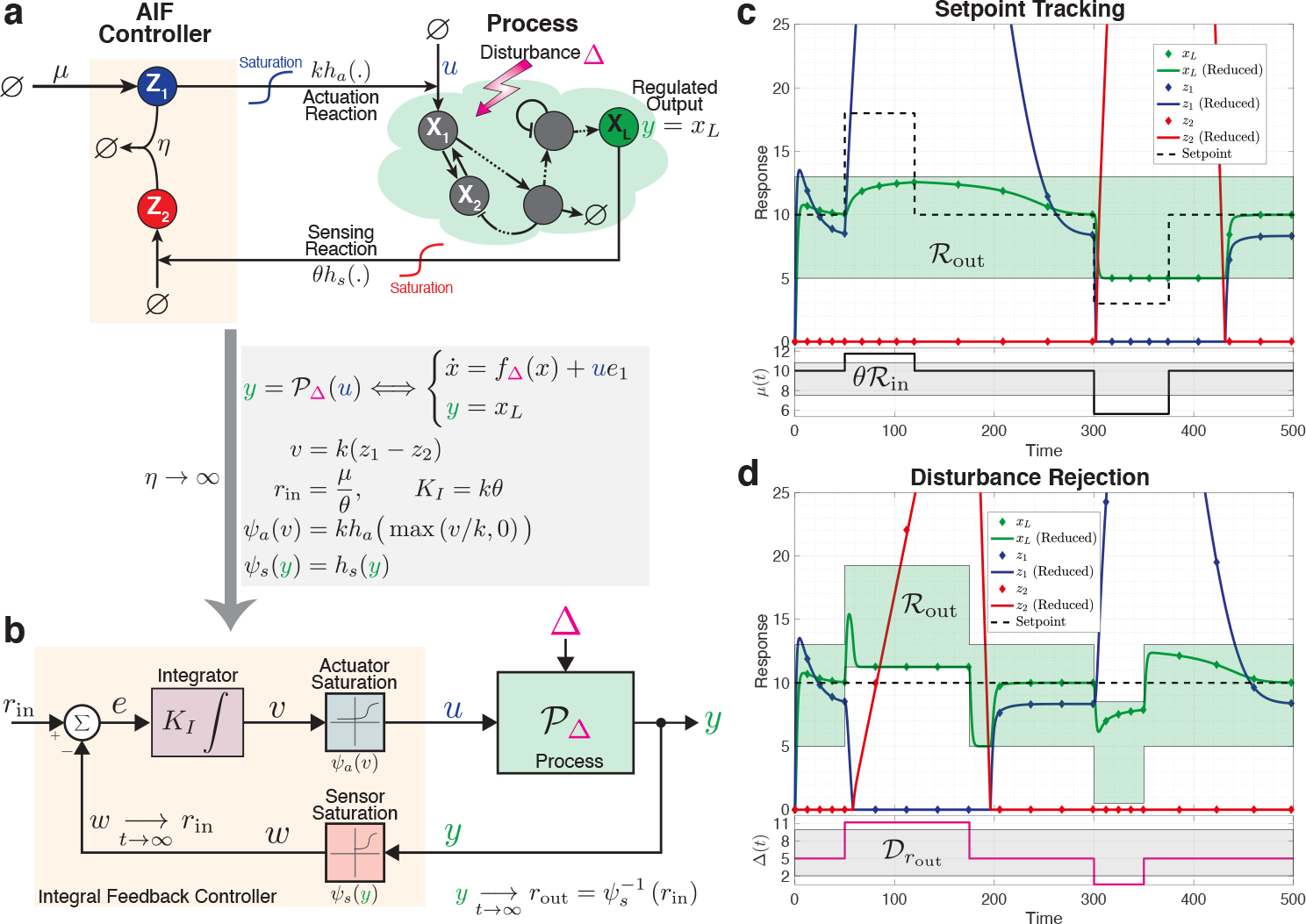
AIF control with actuator and sensor saturation. (a) Closed-loop network. The network structure remains the same as in Fig. 4(a), with the exceptions being the actuation and sensing rates, which are now represented as *u* = *kh*_*a*_(*z*_1_) and *θh*_*s*_(*y*), respectively. Here, *h*_*a*_ and *h*_*s*_ are generally nonlinear, monotonically increasing functions that may be subject to saturation. The input setpoint is designated as *r*_in_ ≜ *µ/θ*, while the output setpoint, *r*_out_, is determined by the sensing function *h*_*s*_ as per (20). (b) Block diagram describing the exact closed-loop dynamics in the limit as *η* → ∞. By defining the integral gain *K*_*I*_ and introducing an intermediate variable *v*, as well as the functions *ψ*_*a*_ and *ψ*_*s*_, we generate this diagram which is similar to that in Fig. 4(b). Here, however, the actuator and sensor saturation modules are replaced by the functions *ψ*_*a*_ and *ψ*_*s*_, respectively. It’s noteworthy that this diagram, once again, mirrors the classical IFC depicted in Fig. 2(c), except for the different forms of actuator and sensor saturation functions. (c) and (d) Integral windup induced by actuator and sensor saturation. For these two simulations, the gene expression model from Fig. 2(b) is used as the process. Panel (c) tests setpoint tracking capabilities when the output setpoint, adjusted via *µ*, becomes transiently inadmissible. In contrast, panel (d) evaluates disturbance rejection properties by maintaining a fixed setpoint while varying the disturbance Δ until it becomes transiently inadmissible. The green shaded areas denote the set of admissible output setpoints ℛ_out_. The gray shaded areas, on the other hand, represent the set of admissible input setpoints (scaled by *θ*) and disturbances in panels (c) and (d), respectively (refer to Example 3 for further details). The simulations reveal that the AIF controller maintains RPA effectively as long as the setpoint and disturbance stay within admissible limits. However, when either transitions to inadmissible levels, RPA is lost and the AIF controller’s performance deteriorates even after returning to admissible levels. This indicates windup manifested by significant accumulation of either *z*_1_ or *z*_2_. Numerical values: *k*_1_ = *γ*_1_ = *γ*_2_ = 1, *η* = 100, *θ* = 15, *κ*_*a*_ = *κ*_*s*_ = 5, *k* = 8. In (c), Δ = 5 is fixed while *µ* is varied; whereas in (d), *µ* = 10 is fixed while Δ is varied.

#### Lemma 2

Consider the closed-loop system depicted in the block diagram of Fig. 5(b), where *ψ*_*a*_ and *ψ*_*s*_ are strictly monotonically increasing functions that may saturate.

For a given desired output setpoint *r*_out_ and a steady-state disturbance or uncertainty 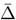, the closed-loop fixed point exists with a feasible supporting input ū ∈ U if and only if:

- The output setpoint is admissible, i.e. *r*_out_ ∈ ℛ_out_.
- The sensor does not saturate at steady-state, i.e. *r*_in_ ∈ range(*ψ*_*s*_).
- The actuator does not saturate at steady-state, i.e. 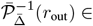 range(*ψ*_*a*_).

Furthermore, the fixed point is given by

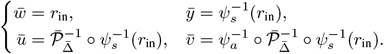

The proof can be found in the Supplementary Information Section S5.

Therefore, as per Corollary 1, for the full model with large *η*, the non-negative closed-loop fixed point will also not exist if any of the conditions of Lemma 2 are violated. However, the closed-loop fixed point may suffer from a negative component rendering it unreachable, and as a result, the dynamics become unstable.

Two numerical simulations are depicted in Fig. 5(c) and (d) where the gene expression model presented in Fig. 2(b) is controlled by the AIF controller that involves actuator and sensor saturation with

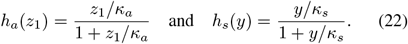

Fig. 5(c) demonstrates setpoint tracking, provided the desired input setpoint, tuned by *µ*, is admissible (i.e., *r*_in_ ∈ ℛ_in_). Similarly, Fig. 5(d) demonstrates disturbance rejection when the disturbance is admissible (i.e. Δ ∈𝒟_*r*_). The sets of admissible input/output setpoints and disturbances, ℛ_in_*/* ℛ_out_ and 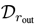, respectively, are calculated in Example 3, by modifying Example 1 to factor in the effects of saturation.

#### Example 3 (Gene Expression with Saturation)

Consider the simple model for gene expression given in Example 1; however, replace the actuation with 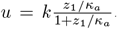. As a result, actuation saturation constrains the feasible inputs to the range 𝕌 = [0, *k*]. The steady-state input/output map remains unchanged as given in Fig. 2(b), resulting in an identical supporting input: 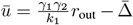.

To ensure the supporting input’s feasibility (0 ≤ ū ≤ *k*), the set of admissible output setpoints is thus given by

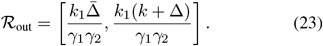

Subsequently, the set of admissible input setpoints *r*_in_ ∈ ℛ_in_ can be mapped from ℛ_out_ using (20), i.e. ℛ_in_ = *h*_*s*_(ℛ_out_). Recalling that *r*_in_ ≜ *µ/θ*, we can outline the gray shaded area in Fig. 5(c) which is given by

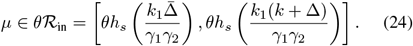

Lastly, for a given output setpoint *r*_out_, the set of admissible disturbances can be computed by determining the disturbances that preserve the feasibility of the supporting input (*ū* ∈ 𝕌), resulting in

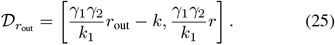

Intuitively, when the (input/output) setpoint exceeds the admissible range, species **Z**_**1**_ accumulates in a futile attempt to increase actuation *u*, which is already saturated. Conversely, when the output setpoint falls below the admissible range, species **Z**_**1**_ essentially tries to become negative to reduce actuation *u* and, as a result, lingers at zero and thus inflicting a buildup of **Z**_**2**_. These observations apply to both Fig. 5(c) and (d).

### F. Anti-Windup Strategies

In this section, we introduce various anti-windup strategies designed to mitigate the unwanted effects of windup. We begin by describing these designs from a phenomenological perspective. We then present their chemical reaction network realizations. Finally, we propose genetic implementations of these anti-windup schemes.

#### 1) Anti-Windup Topologies

The objective here is to present three distinct strategies to alleviate the effects of windup by augmenting the basic AIF motif with extra anti-windup circuitry. These strategies are illustrated as three topologies in Fig. 6(a). Although the three topologies are mechanistically different, they share the same concept: the anti-windup circuitry is only activated to halt further growth when either of the controller species, **Z**_**1**_ or **Z**_**2**_, increases excessively.

**Fig. 6:**
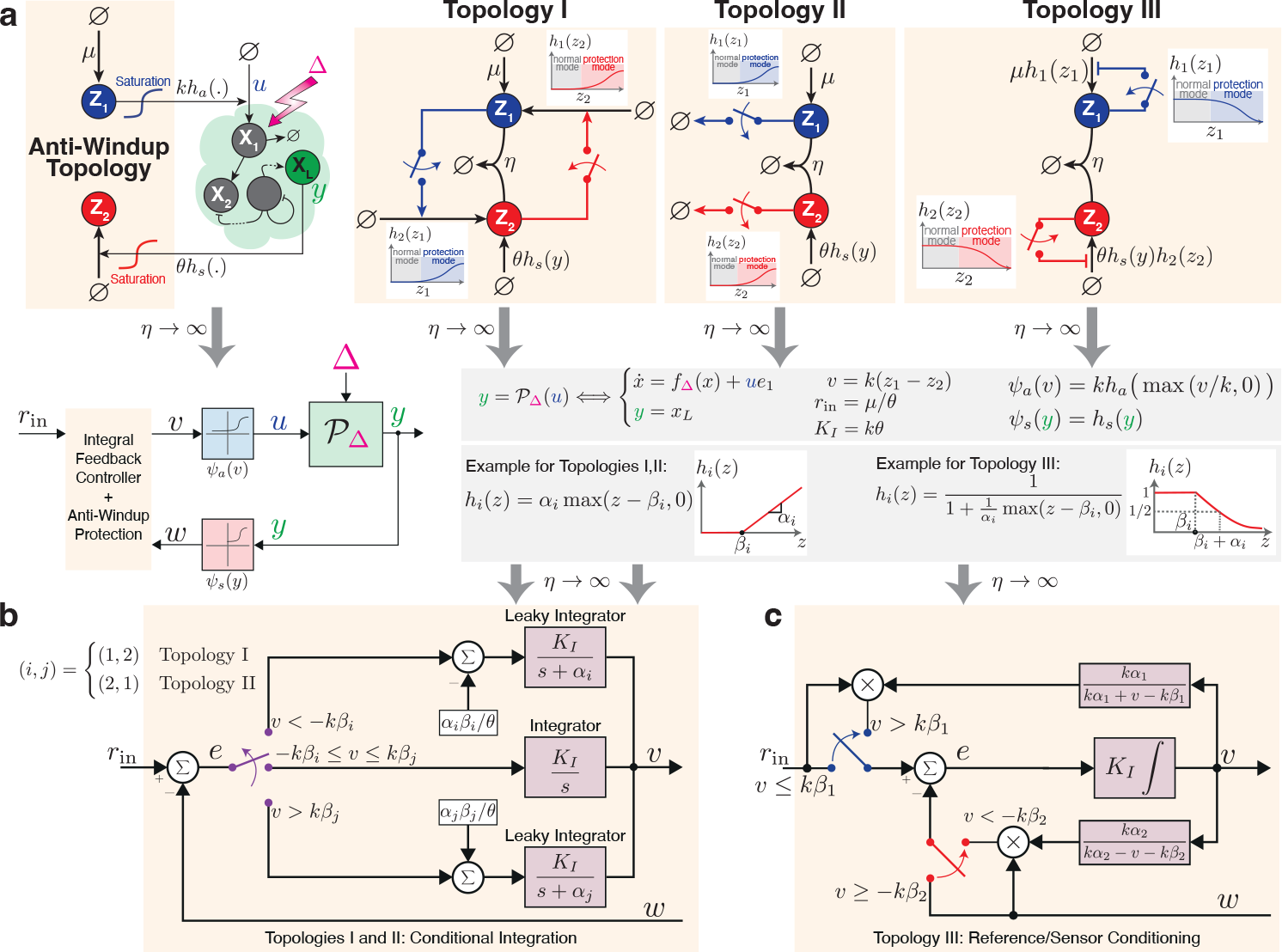
Anti-windup strategies. (a) Three anti-windup topologies. The leftmost diagram depicts a closed-loop network regulated by one of the three distinct anti-windup topologies. In all topologies, the controller operates in normal mode as an AIF controller ensuring RPA, as long as the levels of **Z**_**1**_ and **Z**_**2**_ stay below a defined threshold. However, when either **Z**_**1**_ or **Z**_**2**_ surge beyond this threshold, the anti-windup mechanisms, represented by the red and blue switches, are activated, and the controller transitions into protection mode sacrificing RPA (which is anyway not achievable) for safety. Specifically, Topology I triggers an increased production of **Z**_**1**_ in response to elevated **Z**_**2**_ concentrations, and vice versa. In contrast, Topology II activates the degradation of **Z**_**1**_ and **Z**_**2**_ when their concentrations surpass a defined threshold. Lastly, Topology III reduces the production rates of **Z**_**1**_ and **Z**_**2**_ upon detecting concentrations exceeding a threshold. The switch-like behaviors and corresponding thresholds in these topologies are mathematically represented by the monotonic functions *h*_1_ and *h*_2_. For Topologies I and II, these functions remain essentially zero until the threshold is reached, then increase monotonically. For Topology III, they remain at one and begin decreasing after surpassing the threshold. Concrete mathematical examples of these functions, which can be realized as CRNs, are provided in the lower gray box (refer to Fig. 7 for realization details). In the limit of strong sequestration, introducing the intermediate variable *v*, input setpoint *r*_in_, integral gain *K*_*I*_, and the two functions *ψ*_*a*_ and *ψ*_*s*_, as demonstrated in the top gray box, allows us to generate the block diagram depicted in the bottom left. This diagram portrays the reduced dynamics similar to those presented in Fig. 5(b), but with the addition of the anti-windup dynamics to the AIF controller. (b) A compact block diagram representing Topologies I and II. These topologies employ a “conditional integration” approach, where the controller operates as an integrator when the intermediate variable *v* lies within the safe range [− *kβ*_*i*_, *kβ*_*j*_], and transitions to a “leaky integrator” when operating outside this range to counter windup. The distinction between Topologies I and II is determined by the index pairings (*i, j*), with Topology I associated with (1, 2) and Topology II with (2, 1). (c) A block diagram describing Topology III, which implements a “reference/sensor conditioning” approach. This approach dynamically reduces the input reference or sensor signal when *v* falls outside the safe range [−*kβ*_2_, *kβ*_1_] to prevent windup. Specifically, if *v > kβ*_1_, the input setpoint *r*_in_ is scaled down; conversely, if *v <* −*kβ*_2_, the sensed signal *w* is scaled down.

The anti-windup circuitry for all three topologies are symmetric across **Z**_**1**_ and **Z**_**2**_ (blue and red switches in Fig. 6(a)), thus we will only explain the anti-windup operation on **Z**_**1**_ for brevity. In the case of Topology I, anti-windup is accomplished by having **Z**_**2**_ produce **Z**_**1**_ whenever its concentration reaches a high level. As a result, the produced **Z**_**1**_ species sequester the overabundant **Z**_**2**_ species and thus effectively prevent it from growing any further. Mathematically, this is represented by the monotonically increasing function *h*_1_(*z*_2_), denoting an additional production rate of **Z**_**1**_, which exhibits a switch-like behavior (depicted by the red switch in Fig. 6(a), Topology I). In situations where **Z**_**2**_ concentration remains low, the switch is off, allowing the circuit to function purely as an integral controller. Conversely, when **Z**_**2**_ concentration exceeds a specified threshold, the switch is activated, transitioning the circuit into protection mode. While this mode relinquishes the pure integral control function, it safeguards the system from the detrimental effects of excessive growth. An example of such a function, *h*_1_, is given in the gray box of Fig. 6(a). Here, the production rate remains zero for *z*_2_ ≤ *β*_1_. However, once the threshold *β*_1_ is exceeded, the rate becomes linear in *z*_2_ with a positive slope *α*_1_.

In the case of Topology II, anti-windup is accomplished by having **Z**_**1**_ degrade when its concentration reaches a high level. This strategy is represented by the monotonically increasing function *h*_1_(*z*_1_), denoting a degradation rate of **Z**_**1**_, which exhibits a switch-like behavior (depicted by the blue switch in Fig. 6(a), Topology II).

In Topology III, the anti-windup mechanism is achieved by having **Z**_**1**_ repressing its own production once its concentration surpasses a threshold. Mathematically, this scenario is depicted by the monotonically decreasing function *h*_1_(*z*_1_), which attenuates the maximum production rate *µ* whenever *z*_1_ exceeds the threshold. An example of such a function, *h*_1_, is given in the gray box of Fig. 6(a). Here, the production rate persists at *µ* for *z*_1_ *< β*_1_. However, upon crossing the threshold *β*_1_, the rate decays with **Z**_**1**_. Note that this topology encompasses the anti-windup circuit introduced in [40].

To connect these anti-windup strategies with classical control-theoretic methodologies, we once again look at the asymptotic limit of a large sequestration rate *η*. By invoking the same intermediate variables introduced in Section III-E (and repeated in the gray box of Fig. 6(a) for convenience), we can construct the block diagrams of the controllers illustrated in Fig. 6(b) and (c). The derivations once again rely on Theorem 1 and the details can be found in Section A. Interestingly, both Topologies I and II embody the same control-theoretic principle similar to “conditional integration”; whereas Topology III applies a concept similar to “reference/sensor conditioning” [29], [35], [41].

In the case of conditional integration shown in Fig. 6(b), the controller operates as a pure integral controller, with *r*_in_ being the input setpoint, as long as *v* ≜ *k*(*z*_1_ − *z*_2_) ∈ [− *kβ*_*i*_, *kβ*_*j*_]. However, as soon as *v* departs from this range, the controller behaves like a “leaky integrator”, represented by a transfer function of the form of 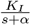. This essentially incorporates a forgetting factor that “forgets” the need to integrate the entire history of the error signal. While leaky integrators introduce a steady-state error, they reciprocate with a stabilizing effect [15]. In fact, the steady-state error can be shown to be proportional to the deviation of *v* from the integrator regime [−*kβ*_*i*_, *kβ*_*j*_]. To see this, we resort to the dynamics of *v* described in the block diagram of Fig. 6(b) that can be explicitly written as

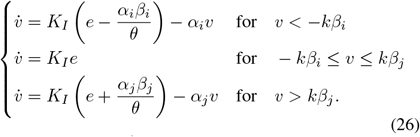

Recalling that *K*_*I*_ ≜ *kθ*, the error at steady-state, when it exists, can be written as

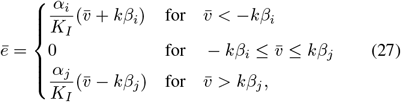

and hence the steady-state error is indeed proportional to the deviation of *v* from the integrator regime.

In the case of reference/sensor conditioning depicted in Fig. 6(c), the controller operates as a pure integral controller, with *r*_in_ being the input setpoint, as long as *v* ≜ *k*(*z*_1_ − *z*_2_) ∈ [− *kβ*_2_, *kβ*_1_]. However, as soon as *v* ventures outside this range, either the reference or sensor signal undergoes a dynamic modification or “conditioning” based on *v*, and thus steering a conditioned version of the error towards zero. This strategy essentially gives up trying to track the desired setpoint when necessary in order to prevent windup. More specifically, if *v > kβ*_1_, then the input reference *r*_in_ is conditioned by reducing it by a factor of *kα*_1_*/*(*kα*_1_ + *v* − *kβ*_1_). Hence, this conditioning becomes more pronounced the further *v* exceeds the integrator operating regime [− *kβ*_2_, *kβ*_1_]. Similarly, if *v < kβ*_2_, then the sensor signal *w* is conditioned by reducing it by a factor of *kα*_2_*/*(*kα*_2_ − *v* − *kβ*_2_). As a result, the conditioning is more significant the further *v* descends from the integrator operating regime.

#### 2) Sequestration-Based Switches

To realize the three anti-windup strategies using CRNs, we first need to realize the switches depicted in Fig. 6(a), which are mathematically represented as the monotonic functions *h*_1_ and *h*_2_. Specifically, we consider the functional forms shown in the gray box of Fig. 6(a) and reproduced in Fig. 7 for convenience. The sequestration networks depicted in Fig. 7(a) and (b) deliver a steady-state response that approximates the desired functions. In fact, this approximation becomes exact in the limit of strong sequestration.

**Fig. 7:**
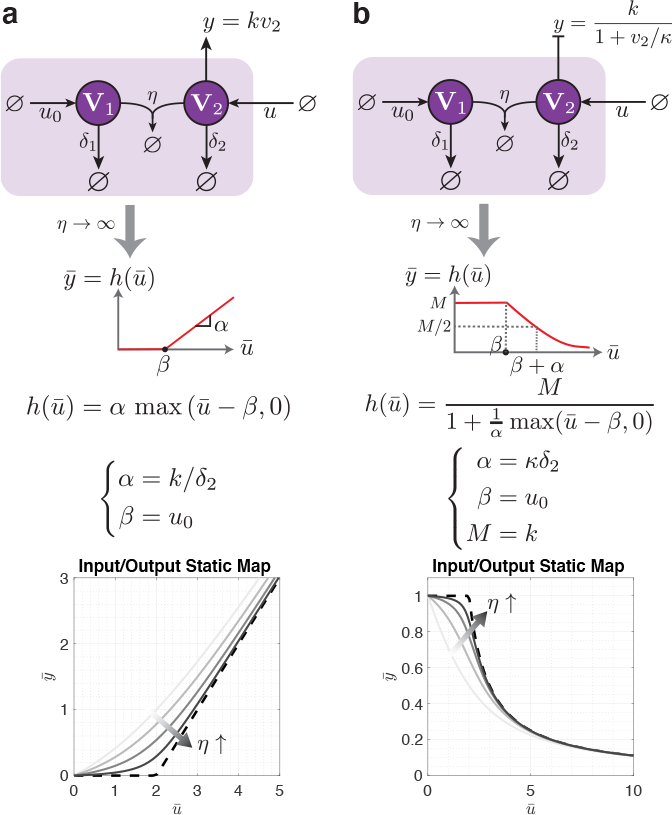
Biomolecular switching using sequestration motifs. The two sequestration networks in (a) and (b) consist two species **V**_**1**_ and **V**_**2**_, which sequester each other at a rate *η* while degrading separately at rates *δ*_1_ and *δ*_2_. The input *u* corresponds to the production rate of **V**_**2**_, and the thresholds are determined by the production rate *u*_0_ of **V**_**1**_. The two networks differ in the role of **V**_**2**_ which acts as an activator in (a) and a repressor in (b). The bottom plots numerically confirm these designs.

The two sequestration networks consist of two species **V**_**1**_ and **V**_**2**_ which sequester each other at a rate of *η*, while separately degrading at rates *δ*_1_ and *δ*_2_, respectively. Here, the input, represented as *u*, signifies the production rate of **V**_**2**_, while the thresholds are determined by the production rate *u*_0_ of **V**_**1**_. The distinguishing feature between the two networks lies in the output: in Fig. 7(a), **V**_**2**_ serves as an activator; conversely, in Fig. 7(b), **V**_**2**_ functions as a repressor. Accordingly, the output is represented by the activation and repression rates, respectively. Once again, Theorems 1 and 2 are invoked here, and the derivations can be found in Section B. These designs are also numerically verified in the plots shown at the bottom of Fig. 7. Notably, decreasing the degradation rates yields a steeper slope leading to an ultrasensitive response [42], [43] – a feature that is not required here for anti-windup.

#### 3) CRN Realizations of Anti-Windup Topologies

Next, we leverage the sequestration-based switches to fully embody the three anti-windup topologies depicted in Fig. 6(a) as CRNs. Fig. 8(a), (b), and (c) each represents a CRN corresponding to anti-windup topologies I, II, and III, respectively.

**Fig. 8:**
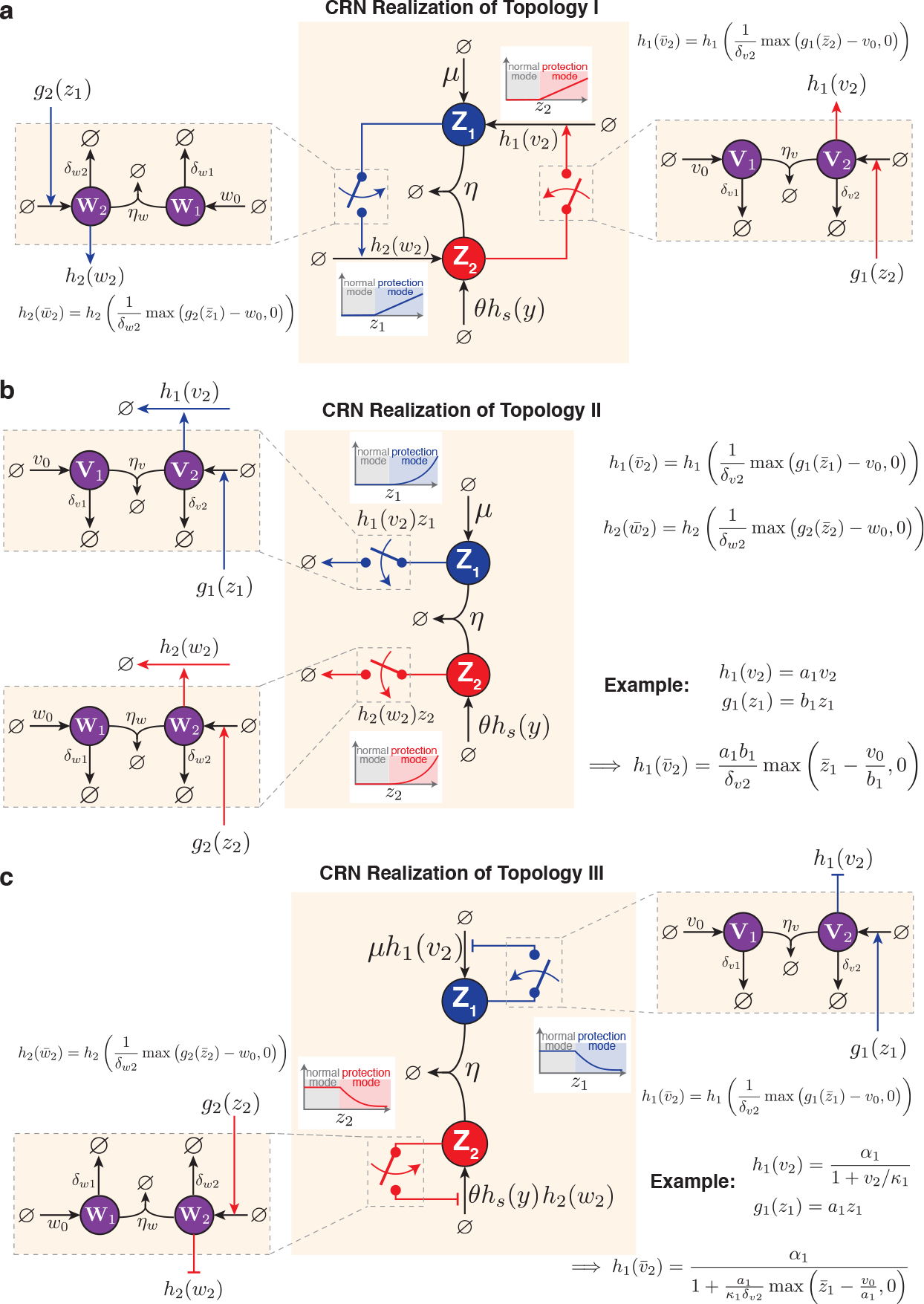
CRN realizations of three anti-windup topologies. (a) Topology I, (b) Topology II, and (c) Topology III. In each topology, sequestration networks, which consist of species (**V**_**1**_,**V**_**2**_) and (**W**_**1**_, **W**_**2**_) that sequester each other, are used to implement switch-like behaviors that intervene to prevent windup when necessary. Topology I employs activator switches and so *h*_1_ and *h*_2_ are monotonically increasing beyond the threshold. Topology II uses a similar mechanism, but the switches catalyze the self-degradations of **Z**_**1**_ and **Z**_**2**_. Here *h*_1_(*v*_2_) and *h*_2_(*w*_2_) represent degradation rates that are monotonically increasing beyond the thresholds. The corresponding propensity functions, however, are given by *h*_1_(*v*_2_)*z*_1_ and *h*_2_(*w*_2_)*z*_2_, respectively. As a result, if *h*_1_ and *h*_2_ are linear functions beyond the threshold, then the corresponding propensities are quadratic. Lastly, Topology III introduces repressor switches, and so *h*_1_(*v*_2_) and *h*_2_(*w*_2_) monotonically decrease beyond the thresholds. Note that while the functions *h*_1_, *h*_2_, *g*_1_, and *g*_2_ are kept intentionally general in this depiction, specific examples are given in panels (b) and (c). In panel (b), the functions are presented in their linear form, while in panel (c), *h*_1_ takes on a Hill form.

Given the symmetry of the anti-windup schemes in **Z**_**1**_ and **Z**_**2**_, we describe the anti-windup operation on **Z**_**1**_ only.

For the CRN realization of Topology I, depicted in Fig. 8(a), the red switch is implemented using the sequestration network from Fig. 7(a). This network consists of two species, **V**_**1**_ and **V**_**2**_, which sequester each other at a rate of *η*_*v*_ and degrade separately at rates *δ*_*v*1_ and *δ*_*v*2_. The production rate of **V**_**2**_, represented as *g*_1_(*z*_2_), serves as the input to the switch and is a monotonically increasing function of *z*_2_. Its output, denoted by *h*_1_(*v*_2_), is a monotonically increasing function of the activator **V**_**2**_, which plays a role in the production rate of **Z**_**1**_. Therefore, with reference to Fig. 7(a) and Section B, assuming a high sequestration rate *η*_*v*_, 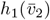 can be written as

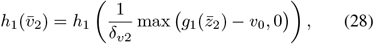

where *v*_0_ is the constant production rate of **V**_**1**_, serving as the threshold. Note that *h*_1_ and *g*_1_ are retained as arbitrary monotonically increasing functions for generality; however, linear representations of these functions would align exactly with the depiction in Fig. 6(a), as shown in the example of Fig. 8(b).

For the CRN realization of Topology II, the blue switch is implemented using a similar sequestration network. The primary differences lie in the production rate of **V**_**2**_, which is now a function of *z*_1_ rather than *z*_2_, and **V**_**2**_ now catalyzes the degradation of **Z**_**1**_ at a rate *h*_1_(*v*_2_). Thus, the degradation propensity of **Z**_**1**_ is represented by *h*_1_(*v*_2_)*z*_1_. With reference to Fig. 7(a) and Section B, assuming a high sequestration rate *η*_*v*_, the degradation propensity at steady-state can be expressed as

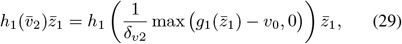

where the propensity now grows quadratically beyond the threshold *v*_0_ (see Fig. 11(b)).

Lastly, for the CRN realization of Topology III, the blue switch is realized using the sequestration network from Fig. 7(b). The main distinction from Topology II lies in **V**_**2**_, which now represses **Z**_**1**_ at a rate *h*_1_(*v*_2_). This rate is a monotonically decreasing function of *v*_2_, for instance, *h*(*v*_2_) = *α*_1_*/*(1 + *v*_2_*/κ*_1_).

#### 4) Genetic Realizations of Anti-Windup Topologies

Genetic realizations of biomolecular integral controllers have already been successfully built and tested in both bacterial and mammalian cells [1]. Therefore, we will not delve into the genetic realization of integral controllers here. Instead, our focus will be on proposing genetic realizations of anti-windup circuitry. The genetic implementations proposed in this study rely on inteins. An intein is a short protein segment that can autocatalytically excise itself from a protein structure and simultaneously rejoin the remaining segments, known as exteins [44]. Split inteins, a subcategory of inteins, are divided into two parts commonly referred to as Int^N^ and Int^C^. When active, these split inteins can heterodimerize and independently carry out protein splicing reactions, which involve the irreversible creation and destruction of peptide bonds in a strict one-to-one stoichiometric ratio. Leveraging their ability to exchange, cleave, or ligate amino acid sequences, split inteins provide the necessary foundation to realize the sequestration reactions that are crucial to the genetic realization of the anti-windup CRNs depicted in Fig. 8. The flexible and small-sized nature of split inteins [45], their existence in numerous orthogonal pairs [46], and their applicability across diverse life forms [47], [48] make them particularly attractive for constructing genetic circuits. In fact, they have previously been employed to build biomolecular integral controllers [9].

Fig. 9(a), (b), and (c) depict three genetic realizations compatible with Topologies I, II, and III from Fig. 8, respectively. Given that the anti-windup circuitry is symmetric with respect to **Z**_**1**_ and **Z**_**2**_, our explanation here will focus on the genetic realization for one of them only. As depicted in Fig. 9(a), the circuit is comprised of two genes that express the proteins **V**_**1**_ and **V**_**2**_. The first gene is driven by a constitutive promoter and includes a degradation tag fused to Int^N^. The second gene, on the other hand, is driven by an inducible promoter activated by **Z**_**2**_. It consists of a DNA-binding domain linked via an Int^C^, and is fused to an activation domain and a degradation tag. Upon expression of the two genes, the corresponding proteins **V**_**1**_ and **V**_**2**_ engage in an intein-splicing reaction and are independently degraded by the proteasome. It’s important to note that prior to splicing, **V**_**2**_ functions as an activator, driving the expression of **Z**_**2**_. However, its function is dismantled when it undergoes the splicing reaction with **V**_**1**_. The threshold of this anti-windup circuit can be adjusted by modifying the strength of the constitutive promoter or its plasmid copy number. It can also be tuned by inducers if the constitutive promoter is replaced by an inducible promoter. In contrast, the anti-windup circuit in Fig. 9(c) is the same as that in Fig. 9(a) but with only two differences: Gene 2 lacks an activation domain and its promoter is activated by **Z**_**1**_ instead of **Z**_**2**_. Consequently, **V**_**2**_ acts as a repressor rather than an activator, making this circuit configuration appropriate for implementing Topology III. Lastly, the anti-windup circuit in Fig. 9(b) is similar to that in Fig. 9(c), except for a single distinction: the DNA-binding domain is replaced by a protease. This modification causes **V**_**2**_ to act as a protease that degrades **Z**_**1**_, rather than repressing its production.

**Fig. 9:**
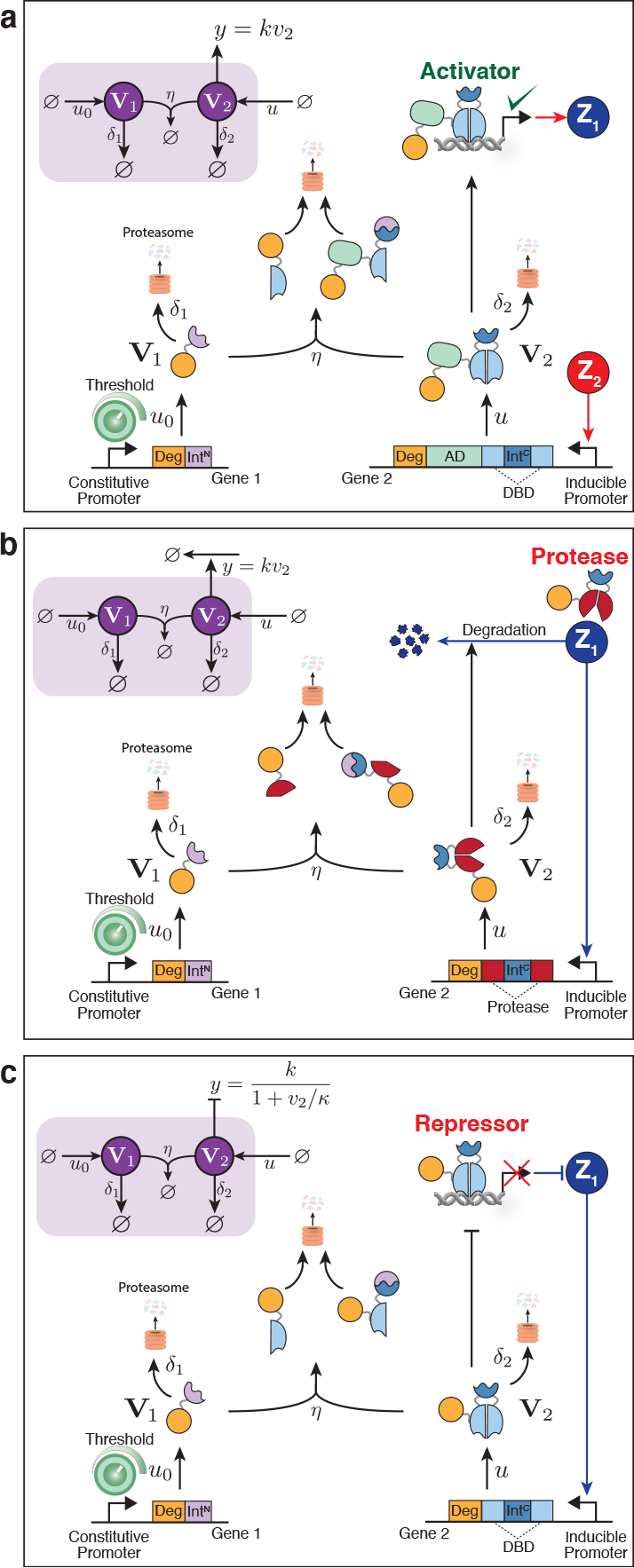
Intein-based genetic realizations of anti-windup topologies. Panels (a), (b), and (c) illustrate the genetic realizations for Topologies I, II, and III, respectively. Each realization is achieved using two genes, one driven by a constitutive promoter and another by an inducible promoter. The proteins expressed by these genes, **V**_**1**_ and **V**_**2**_, engage in an intein-splicing reaction and are subsequently degraded. The function of **V**_**2**_ varies across the topologies acting as an activator in (a), a protease in (b), and a repressor in (c). The strength of the constitutive promoter or plasmid copy number can be manipulated to adjust the anti-windup threshold. AD: activation domain, DBD: DNA binding domain, Int^C^/Int^N^: intein C/N, Deg: degradation tag.

### G. Numerical Simulations

To verify the effectiveness of our proposed anti-windup circuits, we undertake three numerical simulations using three processes of increasing complexity. Fig. 10 depicts the simulations results for Topology I only; however, similar results for Topologies II and III can be found in Supplementary Information, Fig. S1 and S2, respectively.

**Fig. 10:**
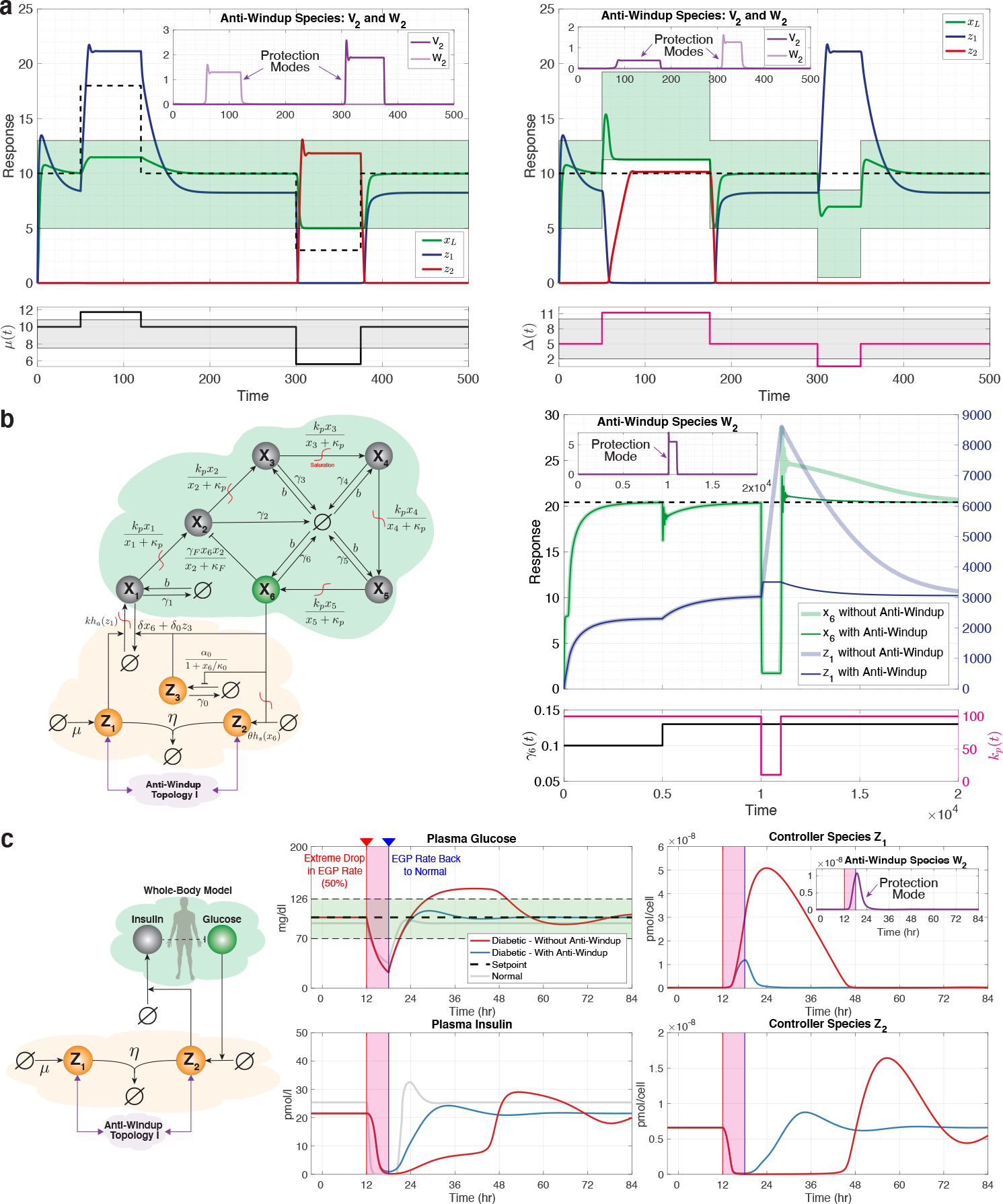
Numerical simulations demonstrating the effectiveness of anti-windup circuitry in three processes of increasing complexity. (a) The first simulation revisits the gene expression process, implementing the CRN realization of Topology I to successfully mitigate windup effects observed in Fig. 5(c) and (d) for both setpoint tracking and disturbance rejection. The parameters of the process and the controller match those in Fig. 5. Anti-windup parameters are set as *η*_*v*_ = *η*_*w*_ = 100, *v*_0_ = 10, *w*_0_ = 20. (b) The second simulation illustrates compatibility with biomolecular PID controllers, where a third-order PID [20] with saturations is employed to regulate a complex process consisting of six species. The plots demonstrate the improvement in dynamic performance of the output **X**_**6**_, resulting from the prevention of **Z**_**1**_ accumulation through the implementation of anti-windup circuitry. Parameter values from [20] are used, while additional parameters are added to introduce saturations, basal expressions, and anti-windup circuitry. The added parameters are as follows: *µ* = 6, *k* = 15, *θ* = 300, *κ*_*a*_ = *κ*_*s*_ = *κ*_*p*_ = 1000, *b* = 0.2, *v*_0_ = *w*_0_ = 3500, and *η*_*v*_ = *η*_*w*_ = *η* = 10^4^. (c) The final simulation employs a mathematical whole-body model of Type I diabetic patients [36], [37], [49] as the controlled process. We test for windup effects under a severe transient disturbance in the endogenous glucose production (EGP) rate. The gray response represents non-diabetic individuals who are not controlled by our circuits, while the blue and red responses represent diabetic patients regulated by the AIF controller with and without anti-windup circuitry, respectively. The plots indicate that glucose levels return smoothly to the healthy range of [70, 126] mg/dl in both non-diabetic individuals and diabetic patients when the AIF controller is supplemented with our anti-windup circuitry. In contrast, without the anti-windup circuitry, the accumulation of **Z**_**1**_ inhibits **Z**_**2**_ from actuating, causing glucose levels to remain elevated beyond the healthy range for an extended period. Parameters are primarily drawn from [10], except for the scaling of *µ* and *θ* by 100 to increase the controller’s speed. Additionally, *η*_*v*_ = *η*_*w*_ = 10^4^ *µ*mol^*−*1^hr^*−*1^, *v*_0_ = 100*w*_0_ = 0.01 *µ*mol/hr are used. Note that the inset plots across all panels show the dynamics of the relevant anti-windup species, which only become active and transition the controller into protection mode when pre-set thresholds are exceeded. Furthermore, we have 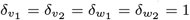, and *h*_1_, *h*_2_, *g*_1_, *g*_2_ from Fig. 8 are all identity functions.

**Fig. 11:**
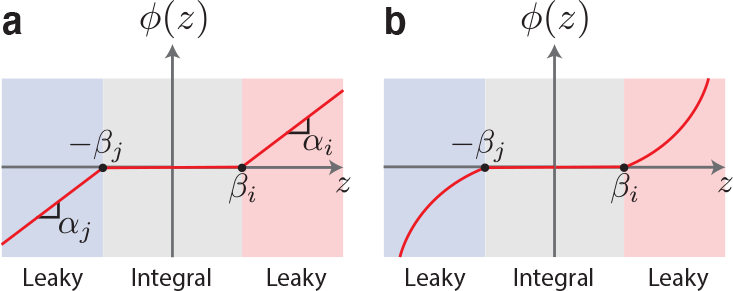
Forgetting functions *ϕ*(*z*). (a) Plot of *ϕ*(*z*) given in (33). (b) Plot of *ϕ*(*z*) given in (39).

The first simulation, shown in Fig. 10(a), revisits the gene expression process outlined in Example 3 – a process that previously demonstrated windup effects as shown in Fig. 5(c) and (d). Here, we utilize the CRN realization of Topology I, shown in Fig. 8(a), to mitigate these windup effects. As depicted in Fig. 10(a), our anti-windup strategy successfully eliminates windup effects in both setpoint tracking (left) and disturbance rejection (right) scenarios. The inset figures track the dynamics of the anti-windup circuits **V**_**2**_ and **W**_**2**_, revealing that they remain dormant until the pre-set thresholds of **Z**_**1**_ and **Z**_**2**_ are surpassed. Upon reaching these thresholds, they become active and shift the controller into a protection mode that sacrifices RPA (which is anyway unattainable) for safety, thereby preventing **Z**_**1**_ and **Z**_**2**_ from reaching excessive levels that would cause windup and deteriorate dynamic performance.

Next, we illustrate that our proposed anti-windup topologies are compatible with biomolecular PID controllers. In this scenario, we consider a more complex process controlled by a third-order PID controller [20], as represented in the closed-loop network of Fig. 10(b). This process consists of six species, **X**_**1**_ through **X**_**6**_, with **X**_**1**_ being the input species and **X**_**6**_ as the regulated output species of interest. Each species **X**_***i***_ degrades at a rate *γ*_*i*_ and catalytically produces **X**_***i*+1**_ for *i* = 1, 2, *· · ·*, 5. In addition, **X**_**6**_ actively degrades **X**_**2**_. This process is taken from [20], [21], but with two modifications: catalytic production reactions are changed to potentially saturating Hill-functions rather than linear functions, and a basal expression rate *b* for all the species is added. The simulation results presented in Fig. 10(b) involve two different disturbances: a persistent but admissible disturbance in *γ*_6_, which the controller successfully rejects, and a transient but inadmissible disturbance in *k*_*p*_ which is the maximum production rate of all species. This intense, albeit brief, disturbance triggers windup as **Z**_**1**_ builds up to significant levels, leading to poor dynamic performance even after the disturbance has passed. However, with the implementation of the anti-windup circuitry, the accumulation of **Z**_**1**_ is circumvented, resulting in superior dynamic performance after. Once again, the inset figure displays the dynamics of the relevant anti-windup species **W**_**2**_, which only activates during windup, transitioning the controller into a protection mode.

In the final simulation example, we utilize a highly complex process pertaining to a mathematical whole-body model of Type I diabetic patients [36], [37], [49]. This model gave rise to the first computer simulator approved by the FDA as an alternative to preclinical trials and animal testing. Note that a prior simulation study demonstrated an effective control of this model using the AIF controller [10], where insulin production is the control action, and blood glucose level is the output required to exhibit RPA. We retain the same AIF closed-loop network and parameter values from [10], with one exception: we increase the parameters *µ* and *θ* by a factor of 100 to speed up the response while maintaining the same setpoint at 100 mg/dl – a value within the healthy range of [70, 126] mg/dl. To test for windup effects, we introduce a severe but transient disturbance in the model. Specifically, the parameter 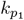 is reduced by 50% for a duration of 6 hours, simulating a dramatic change in endogenous glucose production (EGP), after which the EGP rate returns to normal. The gray curves in Fig. 10(c) correspond to the response of the model of a non-diabetic person not controlled by our circuits. These curves show that insulin secretion abruptly drops to zero to counteract the introduced severe disturbance and successfully returns the glucose level to the healthy range. It’s worth noting that even in a model of a non-diabetic individual, plasma glucose transiently drops to a significantly low level, emphasizing the intensity of the applied disturbance. When we use the AIF controller without the anti-windup circuitry to control the model of a diabetic patient, we get the response depicted in red. The accumulation of **Z**_**1**_ to high levels in response to the inadmissible disturbance is followed by a lengthy recovery time – a clear sign of windup. This leads to a prolonged period of low insulin secretion as the actuator species **Z**_**2**_ remains near zero, resulting in dangerously elevated glucose levels for around 20 hours. However, the implementation of our anti-windup Topology I effectively mitigates this issue. The anti-windup species **W**_**2**_, shown in the inset figure, intervenes during the severe disturbance, preventing **Z**_**1**_ from accumulating and, therefore, avoiding windup. Consequently, glucose levels never overshoot the healthy range.

## IV. Discussion

The phenomenon of windup presents a significant obstacle for any control system that aim to robustly regulate a target variable through integral feedback control. This challenge is especially evident in biomolecular and biomedical contexts where the limitations of actuators, sensors and changes in processes may result in extended periods of poor perfor-mance or even loss of stability. With this in mind, our research focused on designing and evaluating biomolecular anti-windup strategies capable of mitigating windup effects and enhancing the robustness and reliability of biomolecular control systems.

In this study, we introduced three anti-windup topologies specifically adapted for biomolecular control systems that harness the RPA-achieving properties of the AIF controller and its PID extensions. These topologies were carefully designed to account for biomolecular constraints such as positivity, promoter saturation or resource limitations. We initially introduced the topologies using a high-level phenomenological representation involving switches. Subsequently, a sequestration reaction-based CRN was introduced to realize the behavior of the switches. Sequestration motifs are a common fixture in biomolecular designs to realize numerous functionalities including subtraction modules [50], biomolecular input/output linear systems [51], biomolecular integral control [6], biomolecular derivative operators [52]– [54], PID controllers as CRNs [20], [55]–[57], band-pass filters [58], achieving ultra-sensitive response [43], nonlinear activation functions for biomolecular neural networks [59], and stabilization near unstable equilibria [60], among others. This is predominantly attributed to the capability of sequestration reactions to perform comparison operations. In our work, we harnessed this characteristic to realize biomolecular anti-windup strategies, further underlining the essential role of sequestration motifs in biomolecular circuit design.

By leveraging Theorems 1 and 2, we established a connection between these biomolecular topologies and classical control-theoretic concepts. This association was made by calculating transfer functions and block diagrams that are exact in the limit of strong sequestration rates. This approach differs from previous approximation methods that relied on linearizations around the fixed point [20], [21], [54], [57]. It is noteworthy to point out that in [61], a theorem similar to Theorem 1 is presented. In fact, [61] establishes that if there exists some time *T >* 0 before which the concentrations of the sequestration pair do not cross, then simple sequestration networks reduce to subtraction operations for all *t≥ T*. In contrast, Theorem 1 establishes a different model reduction result that is valid for more general networks involving a fast sequestration reaction and under less restrictive conditions. The model reduction results in Theorems 1 and 2 are valid for the dynamics over any compact time interval and for the fixed points, and they require no assumptions except Lipschitz continuity. In a sense, the two approaches complement each other since [61] addresses *t ≥ T* assuming there exists a *T* that satisfies certain conditions; whereas Theorem 1 addresses *t ∈* [0, *T*] for any *T ≥* 0.

The promising potential of our methods to mitigate windup was illustrated both theoretically and through numerical simulations. Our simulations spanned three increasingly complex processes, thereby demonstrating the versatility and robustness of the proposed anti-windup topologies. From a basic gene expression process modeling transcription saturation to a highly intricate whole-body model of a Type I diabetic patient which is FDA approved, our topologies succeeded in preventing windup and maintaining effective control. We also put forth genetic designs capable of implementing the anti-windup strategies, based on inteins which were previously used to successfully build integral controllers. Thanks to their flexibility and small size, inteins hold considerable potential for use in various biomolecular circuits that involve sequestration reactions, such as those in our anti-windup designs.

One primary limitation of the introduced biomolecular anti-windup strategies involves the potential saturation of the anti-windup circuitry itself. As demonstrated in the numerical simulations (particularly the inset figures in Fig. 10), the anti-windup species remain inactive as long as the controller operates within the preset thresholds of normal integral mode. Only when these thresholds are surpassed do the anti-windup species become active to mitigate windup. However, if not properly tuned, this intervention itself may saturate, compromising the effectiveness of the augmented anti-windup circuitry. Nevertheless, even in such a case, the dynamic range of the integrator is expanded compared to a system without anti-windup circuitry, thereby still reducing the impact of windup. Moreover, for the effective implementation of the anti-windup switches depicted in Fig. 6(a) via sequestration reactions, strong sequestrations are paramount. While milder sequestrations can still curtail windup, they introduce steady-state errors even under normal operations (refer to Supplementary Information, Fig. S3). To address this, designers should ensure that sequestration reactions occur more rapidly than other reactions. This is expected to be observed with inteins, where intein-splicing reactions are considerably faster than gene expression processes [62].

Moving forward, the proposed anti-windup topologies represent a promising advancement towards more robust biomolecular control systems. The genetic implementation of these topologies could offer powerful tools for a range of applications, from gene regulation to personalized therapeutics. Moreover, the broader approach of designing safe control systems with consideration for the unique features and constraints of biomolecular processes holds potential to forge new paths in the design of robust and effective biomolecular control systems. A potential avenue for future exploration could involve examining the impacts of windup phenomena and the corresponding anti-windup strategies in a stochastic context.

## V. ACKNOWLEDGMENTS

The authors would like to thank Stanislav Anastassov for the insightful discussions. This research was funded in whole or in part by the Swiss National Science Foundation (SNSF) Grant No. 216505.

## APPENDIX

### A. Anti-Windup Block Diagrams

Consider the closed-loop network depicted in Fig. 6(a) where the controller is given by either Topology I, II or III. We treat each topology separately.

#### Topology I.

The controller dynamics can be expressed as

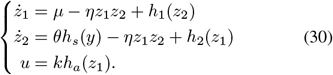

By invoking Theorem 1 as *η* (with *z* ≜ *z*_1_ − *z*_2_, *W*_1_(*x, z*_1_, *z*_2_) ≜ *µ* + *h*_1_(*z*_2_) and *W*_2_(*x, z*_1_, *z*_2_) ≜ *θh*_*s*_(*x*_*L*_) + *h*_2_(*z*_1_)), we obtain the following reduced controller dynamics

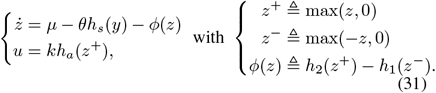

If the functions *h*_1_ and *h*_2_ are given by

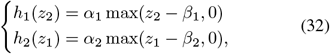

then *ϕ*(*z*) can be expressed as

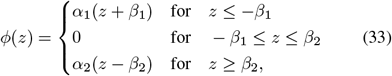

and is plotted in Fig. 11(a). Note that one can think of the function *ϕ* as a “forgetting function” because it causes the integral controller to “forget” the past error signals. Introducing the intermediate variables *v* ≜ *kz, w* ≜ *h*_*s*_(*y*)and *e* ≜ *r*_in_ *− w* and the integral gain *K*_*I*_ ≜ *kθ* yields

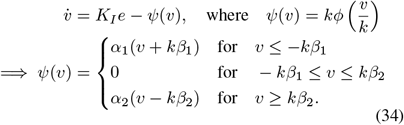

The differential equations for *v* can thus be rewritten as

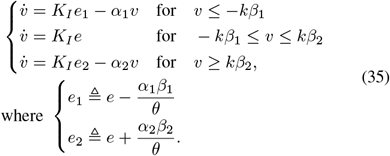

Taking the Laplace transforms to calculate the transfer function from the error *e* to *v* directly gives us the block diagram depicted in Fig. 6(b) with (*i, j*) = (1, 2).

#### Topology II.

The controller dynamics can be expressed as

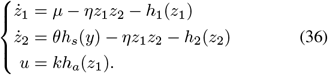

By invoking Theorem 1 as *η* (with *z* ≜ *z*_1_ − *z*_2_, *W*_1_(*x, z*_1_, *z*_2_) ≜ *µ − h*_1_(*z*_1_) and *W*_2_(*x, z*_1_, *z*_2_) ≜ *θh*_*s*_(*x*_*L*_) − *h*_2_(*z*_2_)), we obtain the following reduced con-troller dynamics

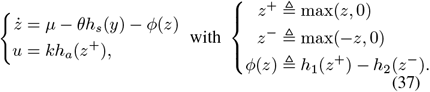

If the functions *h*_1_ and *h*_2_ are also given by (32), then repeating the procedure applied for Topology I will yield an identical block diagram as depicted in Fig. 6(b), but with (*i, j*) = (2, 1). However, if the functions *h*_1_ and *h*_2_ are given by

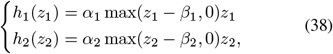

then *ϕ*(*z*) can be expressed as

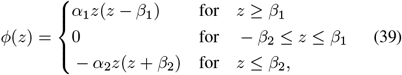

and is plotted in Fig. 11(b). In this case, we do not draw the block diagram because the differential equations for *v* are not piece-wise linear anymore due to the quadratic terms present in *ϕ*(*z*). However, the underlying control-theoretic concept is very similar to that depicted in the block diagram of Fig. 6(b).

#### Topology III.

The controller dynamics can be expressed as

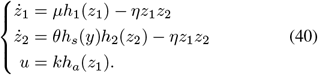

By invoking Theorem 1 as *η* (with *z* ≜ *z*_1_ − *z*_2_, *W*_1_(*x, z*_1_, *z*_2_) ≜ *µh*_1_(*z*_1_) and *W*_2_(*x, z*_1_, *z*_2_) ≜ *θh*_*s*_(*x*_*L*_)*h*_2_(*z*_2_)), we obtain the following reduced controller dynamics

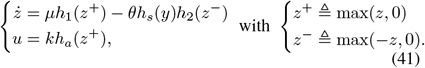

Introducing the intermediate variables *v* ≜ *kz, w* ≜ *h*_*s*_(*y*) and the integral gain *K*_*I*_ ≜ *kθ* yields

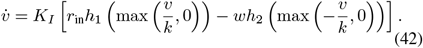

If the functions *h*_1_ and *h*_2_ are given by

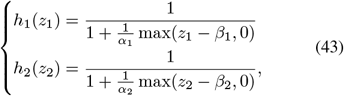

then the differential equations for *v* can be written as

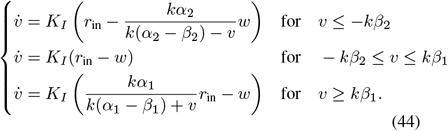

These equations give rise to the block diagram depicted in Fig. 6(c) by defining the error *e* ≜ *r*_in_ *− w*.

#### B. Functional Realizations Via Molecular Sequestration

Consider the two sequestration networks depicted in Fig. 7(a) and (b). The general dynamics for both can be written as

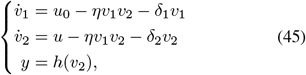

where *u* and *y* denote the input and output of the networks, respectively. The two networks differ by their output functions *h* where we have *h*(*v*_2_) = *kv*_2_ for Fig. 7(a); whereas *h*(*v*_2_) = *k/*(1 +*v*_2_*/κ*) for Fig. 7(b). Note that *h* can take any other functional form. Applying Theorem 1 with *W*_1_(*x, v*_1_, *v*_2_) ≜ *u*_0_ *− δ*_1_*v*_1_ and *W*_2_(*x, v*_1_, *v*_2_) ≜ *u − δ*_2_*v*_2_ yields the following reduced dynamics

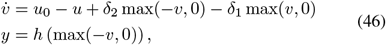

where *v* ≜ *v*_1_ *v*_2_. Hence at steady-state, when it exists, we have

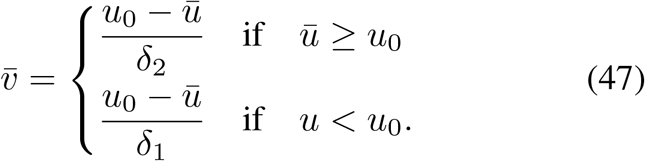

Therefore, invoking Theorem 2 yields the input/output steady-state map of these networks given by

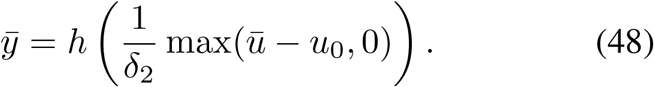

Finally, substituting for the specific form of *h* yields the exact functional forms shown in Fig. 7.

