## Supplementary Information for "Anti-Windup Protection Circuits for Biomolecular Integral Controllers"

#### Contents

**S1 Proof of Lemma 1** 1

**S2 Proof of Theorem 1** 1

**S3 Proof of Theorem 2** 10

**S4 Proof of Corollary 1** 11

**S5 Proof of Lemma 2** 11

**S6 Model Reduction in the Stochastic Setting** 11

**S7 Supplementary Figures** 13

#### S1 Proof of Lemma 1

We first show the sufficiency of these conditions and then their necessity.

*Sufficiency.* Let $r \in [y_{\min}, y_{\max}]$, then the last two equations in (13) imply that $\bar{w} = \bar{y} = r$. If $r \in \mathcal{R}$ and the actuator does not saturate, then the supporting input exists and is given by $\bar{u} = \bar{\mathcal{P}}_{\bar{\Delta}}^{-1}(r) \in [u_{\min}, u_{\max}]$. This implies that $\bar{v}$ exists and is given by $\bar{v} = \bar{u} = \bar{\mathcal{P}}_{\bar{\Delta}}^{-1}(r)$.

*Necessity.* This can be straightforwardly established using the contra-positive, that is, if any of the conditions are not satisfied, then the fixed point does not exist with a feasible supporting input. In fact, if $r \notin \mathcal{R}$ then $\bar{u} \notin \mathbb{U}$. Otherwise, if $\bar{\mathcal{P}}_{\bar{\Delta}}^{-1}(r) \notin [u_{\min}, u_{\max}]$ and/or $r \notin [y_{\min}, y_{\max}]$, then it can be immediately seen from the second and third equations of (13) that $\bar{v}$ and/or $\bar{y}$ do not exist, respectively.

#### S2 Proof of Theorem 1

Consider two deterministic dynamical systems, $\mathcal{S}_\eta$ and $\mathcal{S}$, described by the following set of ODEs and initial conditions

$$\mathcal{S}_\eta : \begin{cases} \dot{x} = F(x, z_1, z_2); & x(0) = x_0 \\ \dot{z}_1 = W_1(x, z_1, z_2) - \eta z_1 z_2; & z_1(0) = a \\ \dot{z}_2 = W_2(x, z_1, z_2) - \eta z_1 z_2; & z_2(0) = b, \end{cases} \qquad \mathcal{S} : \begin{cases} \dot{x} = F(x, z^+, z^-); & x(0) = x_0 \\ \dot{z} = W_1(x, z^+, z^-) - W_2(x, z^+, z^-); & z(0) = \alpha, \end{cases} \tag{1}$$

*Maurice Filo, Ankit Gupta and Mustafa Khammash are with the Department of Biosystems Science and Engineering, ETH Zürich, 4058 Basel, Switzerland.

where $x_0 \in \mathbb{R}_+^L,\ a, b, \eta \in \mathbb{R}_+,\ \alpha \in \mathbb{R}$ and the following notation is adopted

$$z^+ \triangleq \max(z, 0) \quad \text{and} \quad z^- \triangleq \max(-z, 0). \tag{2}$$

The three functions $W_1, W_2 : \mathbb{R}_+^{L+2} \to \mathbb{R}$ and $F : \mathbb{R}_+^{L+2} \to \mathbb{R}^L$ are assumed to be *globally Lipschitz* on their domains and their form is such that the solution of $\mathcal{S}_\eta$ lies in $\mathbb{R}_+^{L+2}$.

We shall prove Theorem 1 in several steps. We first introduce some notation that will be useful throughout the proof. For any two numbers $a$ and $b$, their minimum is denoted by $a \wedge b$. It is immediate that starting from any initial condition $X^{(\eta)}(0) \triangleq (x_0, a, b) \in \mathbb{R}_+^{L+2}$, $\mathcal{S}_\eta$ has a unique solution $X^{(\eta)}(t) \triangleq \left(x^{(\eta)}, z_1^{(\eta)}(t), z_2^{(\eta)}(t)\right)$. Furthermore, it is clear that for large values of $\eta$, the system $\mathcal{S}_\eta$ evolves at two timescales - there is a *slow* timescale of order 1 and a *fast* timescale of order $\eta$. The next lemma examines how the system behaves at the faster timescale.

**Lemma S1** *Let $a, b \in \mathbb{R}_+$ and let $\Psi_t(a, b) \triangleq \big(z_1(t), z_2(t)\big)$ be the solution of the following initial value problem*

$$\begin{cases} \dot{z}_1 = -z_1 z_2; & z_1(0) = a \\ \dot{z}_2 = -z_1 z_2; & z_2(0) = b. \end{cases}$$

*Then for any $t \geq 0$, we can express $\Psi_t(a, b)$ as*

$$\Psi_t(a,b) = \begin{cases} \left(\frac{a(a-b)}{a-be^{-(a-b)t}}, \frac{b(a-b)e^{-(a-b)t}}{a-be^{-(a-b)t}}\right) & \text{if} \quad a > b \\ \left(\frac{a(b-a)e^{-(b-a)t}}{b-ae^{-(b-a)t}}, \frac{b(b-a)}{b-ae^{-(b-a)t}}\right) & \text{if} \quad a < b \\ \left(\frac{a}{1+at}, \frac{a}{1+at}\right) & \text{if} \quad a = b. \end{cases}$$

**Remark S1** *Note that as $t \to \infty$, we get*

$$\Phi(a, b) \triangleq \lim_{t \to \infty} \Psi_t(a, b) = \left((a-b)^+, (a-b)^-\right).$$

*Moreover for any $t \geq 0$, we have*

$$\Psi_t(a,b) - \Phi(a,b) = \begin{cases} \frac{b(a-b)e^{-(a-b)t}}{a-be^{-(a-b)t}}\,(1,1) & \text{if} \quad a > b \\ \frac{a(b-a)e^{-(b-a)t}}{b-ae^{-(b-a)t}}\,(1,1) & \text{if} \quad a < b \\ \frac{a}{1+at}\,(1,1) & \text{if} \quad a = b, \end{cases} \tag{3}$$

*and so the two components of $\Psi_t(a, b) - \Phi(a, b)$ are identical for all $t \geq 0$.*

**Proof.** We shall first prove Lemma S1 in the case $a > b$, hence $\alpha := (a - b) > 0$. Observe that

$$\frac{d(z_1 - z_2)}{dt} = \dot{z}_1 - \dot{z}_2 = 0 \qquad \Longrightarrow \qquad z_1(t) - z_2(t) = z_1(0) - z_2(0) = \alpha.$$

Therefore we can write $z_2(t) = z_1(t) - \alpha$. Substituting this for $z_2$ we obtain

$$\dot{z}_1 = -z_1(z_1 - \alpha), \tag{4}$$

which implies that

$$-\alpha \frac{dz_1}{z_1(z_1 - \alpha)} = \left(\frac{1}{z_1} - \frac{1}{z_1 - \alpha}\right) dz_1 = \alpha dt.$$

Integrating both sides and using $\big(z_1(0), z_2(0)\big) = (a, b)$ we get

$$\log\left(\frac{z_1(t)}{z_1(t) - \alpha}\right) = \log\left(\frac{a}{b}\right) + \alpha t,$$

which upon exponentiation gives us

$$\frac{z_1(t)}{z_1(t)-\alpha} = \left(\frac{a}{b}\right) e^{\alpha t}.$$

Solving for $z_1(t)$ and setting $z_2(t) = z_1(t) - \alpha$ proves the lemma in the case $a > b$. The case $a < b$ follows by symmetry and so we now consider the case $a = b$. It is immediate that in this case $z_1(t) = z_2(t)$ for all $t \geq 0$, and solving the ODE (4) yields

$$z_1(t) = z_2(t) = \frac{a}{1+at}.$$

This completes the proof of this lemma. □

The following lemma establishes that for any given $\eta > 0$, the solution of the system $\mathcal{S}_\eta$ remains bounded over finite time periods.

**Lemma S2** *Let $X^{(\eta)}(t) \triangleq \left(x^{(\eta)}(t), z_1^{(\eta)}(t), z_2^{(\eta)}(t)\right)$ be the solution of system $\mathcal{S}_\eta$. Then, $\forall T > 0$, $\exists M_0 > 0$ such that*

$$\sup_{t\ \in[0,T]} \sup_{\eta} \|X^{(\eta)}(t)\| \leq M_0.$$

**Proof.** From (1) we see that $X^{(\eta)}(t)$ satisfies

$$X^{(\eta)}(t) = X^{(\eta)}(0) + \int_0^t \left(F\left(X^{(\eta)}(s)\right), W_1\left(X^{(\eta)}(s)\right) - \eta z_1^{(\eta)}(s) z_2^{(\eta)}(s), W_2\left(X^{(\eta)}(s)\right) - \eta z_1^{(\eta)}(s) z_2^{(\eta)}(s)\right) ds.$$

Since the components of $X^{(\eta)}(t)$ are always positive the following inequality holds componentwise

$$X^{(\eta)}(t) \leq X^{(\eta)}(0) + \int_0^t \left(F\left(X^{(\eta)}(s)\right), W_1\left(X^{(\eta)}(s)\right), W_2\left(X^{(\eta)}(s)\right)\right) ds.$$

As functions $F, W_1$ and $W_2$ are Lipchitz on $\mathbb{R}_+^{L+2}$, there exists a constant $M \geq 0$ such that

$$\left\|\left(F\left(X^{(\eta)}(t)\right), W_1\left(X^{(\eta)}(t)\right), W_2\left(X^{(\eta)}(t)\right)\right)\right\| \leq M\left\|X^{(\eta)}(t)\right\| + \left\|\Big(F(0), W_1(0), W_2(0)\Big)\right\|, \quad \forall t \in [0,T],$$

where the triangle inequality is employed. This allows us to obtain

$$\begin{aligned}\left\|X^{(\eta)}(t)\right\| &\leq \left\|X^{(\eta)}(0)\right\| + \int_0^t \left(M\left\|X^{(\eta)}(s)\right\| + \left\|\Big(F(0), W_1(0), W_2(0)\Big)\right\|\right) ds \\ &\leq \|(x_0, a, b)\| + \left\|\Big(F(0), W_1(0), W_2(0)\Big)\right\| t + M\int_0^t \left\|X^{(\eta)}(s)\right\| ds.\end{aligned}$$

Applying Gronwall's inequality proves Lemma S2 with $M_0 \triangleq \left(\|(x_0, a, b)\| + \left\|\Big(F(0), W_1(0), W_2(0)\Big)\right\| T\right) e^{MT}$.
□

Next, we examine two specific time instants of particular importance. The first time instant, represented by $\sigma_\eta(\epsilon)$, marks the first moment when $z_1$ and $z_2$ approach within an $\epsilon$-close neighborhood. The second time instant, denoted by $\mu_\eta(\epsilon)$, indicates the first moment when $z_1$ and $z_2$ depart from each other's $2\epsilon$-neighborhood. It should be noted that while $2\epsilon$ is selected here for simplicity, any value $k\epsilon$ could be chosen, provided $k > 1$.

**Proposition S1** *Let* $X^{(\eta)}(t) \triangleq \left(x^{(\eta)}(t), z_1^{(\eta)}(t), z_2^{(\eta)}(t)\right)$ *be the solution of system* $\mathcal{S}_\eta$*, but with initial condition* $X^{(\eta)}(0)$ *satisfying* $\lim_{\eta\to\infty} X^{(\eta)}(0) = (x_0, \epsilon, 0)$ *for some* $x_0 \in \mathbb{R}_+^L$ *and* $\epsilon > 0$*. Furthermore, let* $\big(x(t), z(t)\big)$ *be the solution of system* $\mathcal{S}$*, but with initial condition* $\big(x(0), z(0)\big) = (x_0, \epsilon)$*. Define*

$$\mu_\eta(\epsilon) \triangleq \inf\left\{t \geq 0 : \left|z_1^{(\eta)}(t) - z_2^{(\eta)}(t)\right| \geq 2\epsilon\right\}.$$

*Then we have*

$$\liminf_{\eta\to\infty} \mu_\eta(\epsilon) > 0, \tag{5}$$

*and for any* $T > 0$*, there exists two constants* $C_1, C_2 > 0$ *such that*

$$\limsup_{\eta\to\infty} \frac{\sup_{t\in[0,\mu_\eta(\epsilon)\wedge T]} \left\|\left(x^{(\eta)}(t), z_1^{(\eta)}(t) - z_2^{(\eta)}(t)\right) - \big(x(t), z(t)\big)\right\|}{(\mu_\eta(\epsilon)\wedge T)e^{C_1(\mu_\eta(\epsilon)\wedge T)}} \leq C_1\epsilon \tag{6}$$

$$\limsup_{\eta\to\infty} \frac{\int_0^{\mu_\eta(\epsilon)\wedge T} \left\|X^{(\eta)}(t) - \big(x(t), z^+(t), z^-(t)\big)\right\| dt}{\mu_\eta(\epsilon)\wedge T} \leq C_2\epsilon. \tag{7}$$

**Remark S2** *By symmetry, the proposition also holds with initial conditions* $\lim_{\eta\to\infty} X^{(\eta)}(0) = (x_0, 0, \epsilon)$ *for* $\mathcal{S}_\eta$ *and* $\big(x(0), z(0)\big) = (x_0, -\epsilon)$ *for* $\mathcal{S}$*. It will be clear in the proof that the proposition also holds if* $\lim_{\eta\to\infty} X^{(\eta)}(0) = (x_0, a, b) \in \mathbb{R}_+^{L+2}$ *and* $\big(x(t), z(t)\big) = (x_0, a - b)$*, where either* $a \leq \epsilon$ *and* $b = 0$ *or* $a = 0$ *and* $b \leq \epsilon$*.*

**Proof.** Without loss of generality we can assume that $X^{(\eta)}(0) = (x_0, \epsilon, 0)$ for each $\eta$. Observe that

$$\frac{d}{dt}\left(z_1^{(\eta)}(t) - z_2^{(\eta)}(t)\right) = W_1\left(X^{(\eta)}(t)\right) - W_2\left(X^{(\eta)}(t)\right),$$

and since $z_1^{(\eta)}(0) - z_2^{(\eta)}(0) = \epsilon$, the definition of $\mu_\eta(\epsilon)$ implies that

$$2\epsilon = \left|\epsilon + \int_0^{\mu_\eta(\epsilon)} \left[W_1\left(X^{(\eta)}(t)\right) - W_2\left(X^{(\eta)}(t)\right)\right] dt\right|.$$

Using Lemma S2 and the Lipchitz conditions on functions $W_1$ and $W_2$, we can find a constant $W_0 > 0$ satisfying

$$\sup_{t\in[0,T]} \sup_\eta \left|W_1\left(X^{(\eta)}(t)\right) - W_2\left(X^{(\eta)}(t)\right)\right| \leq W_0.$$

This ensures that $\mu_\eta(\epsilon) \geq \epsilon/W_0$ and proves (5). Next, define a function $\bar{F}$ by transforming the function $F$ as follows

$$\bar{F}(x, z_1, z_2) \triangleq F\left(x, \frac{z_1 + z_2}{2}, \frac{z_2 - z_1}{2}\right).$$

Also define the functions $\bar{W}_1$ and $\bar{W}_2$ by similarly transforming functions $W_1$ and $W_2$, respectively. Let

$$\chi^{(\eta)}(t) \triangleq \left(x^{(\eta)}(t), v^{(\eta)}(t), w^{(\eta)}(t)\right), \quad \text{where} \begin{cases} v^{(\eta)}(t) \triangleq z_1^{(\eta)}(t) - z_2^{(\eta)}(t) \\ w^{(\eta)}(t) \triangleq z_1^{(\eta)}(t) + z_2^{(\eta)}(t). \end{cases}$$

One can see that $\chi^{(\eta)}(t)$ satisfies the following system of ODEs

$$\begin{cases} \dot{x} = \bar{F}(x, v, w); & x(0) = x_0 \\ \dot{v} = \bar{W}_1(x, v, w) - \bar{W}_2(x, v, w); & v(0) = \epsilon \\ \dot{w} = \bar{W}_1(x, v, w) + \bar{W}_2(x, v, w) - \frac{1}{2}\eta(w^2 - v^2); & w(0) = \epsilon. \end{cases}$$

Moreover $\big(x(t), z(t)\big)$ satisfies the following system of ODEs

$$\begin{cases} \dot{x} = \bar{F}(x, z, |z|); & x(0) = x_0 \\ \dot{z} = \bar{W}_1(x, z, |z|) - \bar{W}_2(x, z, |z|); & z(0) = \epsilon. \end{cases}$$

Note that $v^{(\eta)}(0) = \epsilon$, and we can express $\mu_\eta(\epsilon)$ as

$$\mu_\eta(\epsilon) = \inf\left\{t \geq 0 : \left|v^{(\eta)}(t)\right| \geq 2\epsilon\right\}.$$

As before, we can find a constant $W_0$ such that for any $\eta$ we have

$$\sup_{t\in[0,T]} \left|W_1\left(X^{(\eta)}(t)\right) + W_2\left(X^{(\eta)}(t)\right)\right| \leq W_0.$$

This ensures that in the time interval $[0, \mu_\eta(\epsilon))$, the differential inequality

$$\dot{w} \leq W_0 + 2\eta\epsilon^2 - \frac{1}{2}\eta w^2; \quad w(0) = \epsilon,$$

is satisfied by $w^{(\eta)}(t)$. This means that in this time interval, we have $w^{(\eta)}(t) \leq \gamma(t)$ where $\gamma(t)$ satisfies

$$\dot{\gamma} = W_0 + 2\eta\epsilon^2 - \frac{1}{2}\eta\gamma^2; \quad \gamma(0) = \epsilon.$$

Solving this initial value problem, we obtain

$$\gamma(t) = c\left[\frac{(\epsilon + c) + (\epsilon - c)e^{-\eta ct}}{(\epsilon + c) - (\epsilon - c)e^{-\eta ct}}\right], \qquad \text{with } c = 2\epsilon\sqrt{1 + \frac{W_0}{2\epsilon^2\eta}}.$$

This shows that

$$\limsup_{\eta\to\infty} \frac{1}{\mu_\eta(\epsilon) \wedge T} \int_0^{\mu_\eta(\epsilon)\wedge T} |w^{(\eta)}(t)|dt \leq 2\epsilon. \tag{8}$$

One can see that

$$\Big(x^{(\eta)}(t), v^{(\eta)}(t)\Big) - \Big(x(t), z(t)\Big) =$$

$$\int_0^t \left(\bar{F}\left(x^{(\eta)}(s), v^{(\eta)}(s), w^{(\eta)}(s)\right) - \bar{F}\Big(x(s), z(s), |z(s)|\Big), \bar{W}\left(x^{(\eta)}(s), v^{(\eta)}(s), w^{(\eta)}(s)\right) - \bar{W}\Big(x(s), z(s), |z(s)|\Big)\right)ds,$$

where $\bar{W}(x, v, w) = \bar{W}_1(x, v, w) - \bar{W}_2(x, v, w)$. This integral relation along with Lipchitz conditions on functions $\bar{F}$ and $\bar{W}$ implies that there exists a constant $M > 0$ such that

$$\left\|\Big(x^{(\eta)}(t), v^{(\eta)}(t)\Big) - \Big(x(t), z(t)\Big)\right\| \leq M\int_0^t \left(\left\|x^{(\eta)}(s) - x(s)\right\| + \left|v^{(\eta)}(s) - z(s)\right| + \left|w^{(\eta)}(s) - |z(s)|\right|\right)ds.$$

But $\left|w^{(\eta)}(s) - |z(s)|\right| \leq \left|w^{(\eta)}(s) - |v^{(\eta)}(s)|\right| + \left||v^{(\eta)}(s)| - |z(s)|\right| \leq \left|w^{(\eta)}(s)\right| + \left|v^{(\eta)}(s) - z(s)\right|$, because $w^{(\eta)}(s) \geq \left|v^{(\eta)}(s)\right|$ and $\left||a| - |b|\right| \leq \left|a - b\right|$ for any $a, b \in \mathbb{R}$. For any $t \in [0, \mu_\eta(\epsilon) \wedge T]$ and large $\eta$ this allows us to write

$$\begin{aligned}\left\|\Big(x^{(\eta)}(t), v^{(\eta)}(t)\Big) - \Big(x(t), z(t)\Big)\right\| &\leq M\int_0^t \left(\left\|x^{(\eta)}(s) - x(s)\right\| + \left|v^{(\eta)}(s) - z(s)\right| + \left|w^{(\eta)}(s) - |z(s)|\right|\right)ds \\ &\leq M\int_0^t \left|w^{(\eta)}(s)\right|ds + 2M\int_0^t \left\|\Big(x^{(\eta)}(s), v^{(\eta)}(s)\Big) - \Big(x(s), z(s)\Big)\right\|ds \\ &\leq 2M\epsilon(\mu_\eta(\epsilon) \wedge T) + 2M\int_0^t \left\|\Big(x^{(\eta)}(s), v^{(\eta)}(s)\Big) - \Big(x(s), z(s)\Big)\right\|ds,\end{aligned}$$

where the last inequality follows from (8). Using Gronwall's inequality we can conclude that with $C_1 = 2M$, we have

$$\limsup_{\eta\to\infty} \sup_{t\in[0,\mu_\eta(\epsilon)\wedge T]} \frac{\left\| \left(x^{(\eta)}(t), v^{(\eta)}(t)\right) - \left(x(t), z(t)\right) \right\|}{(\mu_\eta(\epsilon)\wedge T)e^{C_1\mu_\eta(\epsilon)\wedge T}} \leq C_1\epsilon, \tag{9}$$

which proves (6). Combining this with (8) and $\left|w^{(\eta)}(s) - |z(s)|\right| \leq \left|w^{(\eta)}(s)\right| + \left|v^{(\eta)}(s) - z(s)\right|$ we can easily show (7). This completes the proof of this proposition. □

Let the functions $\Psi_t, \Phi : \mathbb{R}_+^2 \to \mathbb{R}_+^2$ be as in Lemma S1 and Remark S1, respectively. We extend these functions to functions from $\mathbb{R}_+^2$ to $\mathbb{R}_+^{L+2}$ by defining

$$\widehat{\Psi}_t(x, a, b) \triangleq (x, \Psi_t(a, b)) \qquad \text{and} \qquad \widehat{\Phi}(x, a, b) \triangleq (x, \Phi(a, b)).$$

Recall that $X^{(\eta)}(t) = \left(x^{(\eta)}(t), z_1^{(\eta)}(t), z_2^{(\eta)}(t)\right)$ is a solution to the system $\mathcal{S}_\eta$, and define another process $Y^{(\eta)}(t)$ as

$$Y^{(\eta)}(t) \triangleq X^{(\eta)}(t) - \left[\widehat{\Psi}_{\eta t}\left(X^{(\eta)}(0)\right) - \widehat{\Phi}\left(X^{(\eta)}(0)\right)\right]. \tag{10}$$

Note that $Y^{(\eta)}(t)$ has the form $Y^{(\eta)}(t) = \left(x^{(\eta)}(t), y_1^{(\eta)}(t), y_2^{(\eta)}(t)\right)$, where

$$\left(y_1^{(\eta)}(t), y_2^{(\eta)}(t)\right) = \left(z_1^{(\eta)}(t), z_2^{(\eta)}(t)\right) - \left[\Psi_{\eta t}\left(z_1^{(\eta)}(0), z_2^{(\eta)}(0)\right) - \Phi\left(z_1^{(\eta)}(0), z_2^{(\eta)}(0)\right)\right]. \tag{11}$$

For time $t$ bounded away from 0, $\widehat{\Psi}_{\eta t}\left(X^{(\eta)}(0)\right) - \widehat{\Phi}\left(X^{(\eta)}(0)\right)$ would be close to $(0, 0, 0)$; whereas for $t$ near 0, $X^{(\eta)}(t) - \widehat{\Psi}_{\eta t}\left(X^{(\eta)}(0)\right)$ would be close to $(0, 0, 0)$. As it turns out, process $Y^{(\eta)}(t)$ has better convergence properties (as $\eta \to \infty$) in comparison to process $X^{(\eta)}(t)$. We shall use the results in Katzenberger [1], to prove the following proposition which will play a key role in the proof of Theorem 1.

**Proposition S2** *Suppose that* $\lim_{\eta\to\infty}\left(x^{(\eta)}(0), z_1^{(\eta)}(0) - z_2^{(\eta)}(0)\right) = (x_0, \alpha)$ *for some* $(x_0, \alpha) \in \mathbb{R}_+^L \times \mathbb{R}$, *and let* $\left(x(t), z(t)\right)$ *be the solution of system* $\mathcal{S}$ *with initial condition* $(x_0, \alpha)$. *Fix a small* $\epsilon > 0$, *and define* $\sigma_\eta(\epsilon)$ *as*

$$\sigma_\eta(\epsilon) \triangleq \inf\left\{t \geq 0 : \left|z_1^{(\eta)}(t) - z_2^{(\eta)}(t)\right| \leq \epsilon\right\}. \tag{12}$$

*Then we have the following*

*(A)* $\lim_{\eta\to\infty} \sigma_\eta(\epsilon) = \sigma(\epsilon) \triangleq \inf\{t \geq 0 : |z(t)| \leq \epsilon\}$.

*(B) If* $\sigma(\epsilon) \in (0, \infty)$ *then*

$$\left(z^+\left(\sigma(\epsilon)\right), z^-\left(\sigma(\epsilon)\right)\right) = \begin{cases} (\epsilon, 0) & \text{if} \quad \alpha > 0 \\ (0, \epsilon) & \text{if} \quad \alpha < 0. \end{cases}$$

*and for any* $T > 0$

$$\limsup_{\eta\to\infty} \sup_{t\in[\delta,\sigma_\eta(\epsilon)\wedge T]} \left\| X^{(\eta)}(t) - \left(x(t), z^+(t), z^-(t)\right) \right\| = 0, \tag{13}$$

*where* $\delta$ *is any positive number less that* $\sigma(\epsilon)$.

**Proof.** Without loss of generality we can assume that there is a $\theta \in \mathbb{R}_+$ such that $X^{(\eta)}(0) = (x_0, \theta + \alpha, \theta)$ for all $\eta$. Clearly the assertions of this proposition become trivial when $|\alpha| \leq \epsilon$, since $\sigma(\epsilon) = 0$, and so we can assume that $|\alpha| > \epsilon$.

We shall prove the proposition under the assumption $\alpha > \epsilon$. The other case $\alpha < -\epsilon$ follows by symmetry. We first consider the case $\theta > 0$. Define $\Gamma = \{(x, z, 0) \in \mathbb{R}_+^{L+2} : x \in \mathbb{R}_+^L \text{ and } z > 0\}$ and $U_\Gamma = \{(x, z_1, z_2) \in \mathbb{R}_+^{L+2} : x \in \mathbb{R}_+^L \text{ and } z_1 > z_2 \geq 0\}$. Observe that $U_\Gamma$ is an open set in $\mathbb{R}_+^{L+2}$ containing $\Gamma$. Moreover $\Gamma$ is an $(L-1)$-dimensional continuous manifold and the function $\widehat{\Phi}$ is continuously differentiable on $\Gamma$. In what follows, we denote the vector or matrix of zeros by $\mathbf{0}$. Letting $G(x, z_1, z_2) \triangleq (\mathbf{0}, -z_1 z_2, -z_1 z_2)$ and $W(x, z_1, z_2) \triangleq \Big(F(x, z_1, z_2), W_1(x, z_1, z_2), W_2(x, z_1, z_2)\Big)$, we can see that $X^{(\eta)}(t)$ satisfies

$$\dot{X}^{(\eta)} = W\left(X^{(\eta)}(t)\right) + \eta G\left(X^{(\eta)}(t)\right); \qquad X^{(\eta)}(0) = (x_0, \theta + \alpha, \theta).$$

Also at any $(x, z, 0) \in \Gamma$, the Jacobian matrix $\partial G(x, z, 0)$ of function $G$ can be computed as

$$\partial G(x, z, 0) = z \begin{bmatrix} \mathbf{0} & \mathbf{0} & \mathbf{0} \\ \mathbf{0} & -1 & 0 \\ \mathbf{0} & -1 & 0 \end{bmatrix},$$

and it has exactly one negative eigenvalue. For any $t \in [0, \sigma_\eta(\epsilon)]$, we have $Y^{(\eta)}(t) \in U_\Gamma$ for large $\eta$ due to (3). Let $\widetilde{Y}^{(\eta)}(t)$ denote the trajectory of $Y^{(\eta)}(t)$ stopped at time $\sigma_\eta(\epsilon)$, i.e.

$$\widetilde{Y}^{(\eta)}(t) = Y^{(\eta)}(t \wedge \sigma_\eta(\epsilon)) \qquad \text{for} \qquad t \geq 0.$$

From Theorem 6.3 in [1] we can conclude that the sequence $(\widetilde{Y}^{(\eta)}, \sigma_\eta(\epsilon))$ converges in the Skorohod topology on $D_{\mathbb{R}_+^2}[0, \infty) \times [0, \infty]$ to $(\widetilde{Y}, \sigma(\epsilon))$, where

$$\sigma(\epsilon) = \inf\left\{t \geq 0 : |\widetilde{y}_1(t) - \widetilde{y}_2(t)| < \frac{\epsilon}{2}\right\}, \tag{14}$$

and the trajectory $\widetilde{Y}(t) = \Big(\widetilde{x}(t), \widetilde{y}_1(t), \widetilde{y}_2(t)\Big)$ lies in the set $\Gamma$ and satisfies

$$\widetilde{Y}(t) = \widetilde{Y}(0) + \int_0^{t \wedge \sigma(\epsilon)} \partial\widehat{\Phi}\Big(\widetilde{Y}(s)\Big) W\Big(\widetilde{Y}(s)\Big) ds. \tag{15}$$

However $\widehat{\Phi}(x, z_1, z_2) = (x, z_1 - z_2, 0)$ on $U_\Gamma$, and hence the Jacobian matrix is simply

$$\partial\Phi(x, z, 0) = \begin{bmatrix} \mathbf{I} & \mathbf{0} & \mathbf{0} \\ \mathbf{0} & 1 & -1 \\ \mathbf{0} & 0 & 0 \end{bmatrix},$$

for any $(x, z, 0) \in \Gamma$, where $\mathbf{I}$ is the $L \times L$ identity matrix. Hence we can rewrite equation (15) for $\widetilde{Y}(t) = \big(\widetilde{x}(t), \widetilde{y}_1(t), \widetilde{y}_2(t)\big)$ as

$$\begin{aligned} \widetilde{x}(t) &= \widetilde{x}(0) + \int_0^{t \wedge \sigma(\epsilon)} F\big(\widetilde{x}(s), \widetilde{y}_1(s), \widetilde{y}_2(s)\big) ds \\ \widetilde{y}_1(t) &= \widetilde{y}_1(0) + \int_0^{t \wedge \sigma(\epsilon)} \Big(W_1\big(\widetilde{x}(s), \widetilde{y}_1(s), \widetilde{y}_2(s)\big) - W_2\big(\widetilde{x}(s), \widetilde{y}_1(s), \widetilde{y}_2(s)\big)\Big) ds \\ \text{and} \quad \widetilde{y}_2(t) &= 0. \end{aligned}$$

Since $X^{(\eta)}(0) = (x_0, \theta + \alpha, \theta)$, we see that $\widetilde{Y}(0) = \big(\widetilde{x}(0), \widetilde{y}_1(0), \widetilde{y}_2(0)\big) = \widehat{\Phi}(x_0, \theta + \alpha, \theta) = (x_0, \alpha, 0)$. Hence $\widetilde{x}(0) = x_0$ and $\widetilde{y}_1(0) = \alpha$, and so we have $\big(\widetilde{x}(t), \widetilde{y}_1(t)\big) = \big(x(t), z(t)\big)$ in the time interval $[0, \sigma(\epsilon)]$, where $(x(t), z(t))$ is as in the statement of this proposition. As $\widetilde{y}_2(t) = 0$ until time $\sigma(\epsilon)$, we can express (14) as

$$\sigma(\epsilon) = \inf\left\{t \geq 0 : 0 < z(t) < \epsilon\right\}. \tag{16}$$

This proves part (A) of the proposition.

Suppose $\sigma(\epsilon) < \infty$ and since $\alpha > \epsilon$ we also have $\sigma(\epsilon) > 0$. Due to continuity of the trajectories $z(\sigma(\epsilon)) = \epsilon$. These facts along with the convergence $\left(\widetilde{Y}^{(\eta)}, \sigma_\eta(\epsilon)\right) \longrightarrow \left(\widetilde{Y}, \sigma(\epsilon)\right)$ in the Skorohod topology and (3), imply that

$$\limsup_{\eta\to\infty} \sup_{t\in[\delta,\sigma_\eta(\epsilon)\wedge T]} \left\| X^{(\eta)}(t) - \big(x(t), z^+(t), z^-(t)\big) \right\| = \limsup_{\eta\to\infty} \sup_{t\in[\delta,\sigma_\eta(\epsilon)\wedge T]} \left\| Y^{(\eta)}(t) - \big(x(t), z^+(t), z^-(t)\big) \right\| = 0,$$

thereby proving part (B).

We now come to the case $\theta = z_2^{(\eta)}(0) = 0$. In this case we can set $Y^{(\eta)} = Z^{(\eta)}$ and use Theorem 6.2 in [1] to conclude the above convergence in the Skorohod topology. The rest of the arguments go through as before. This concludes the proof of this proposition. □

**Lemma S3** *Fix a $T > 0$, $\epsilon_0 > 0$ and an $\epsilon \in (0, \epsilon_0)$. Suppose for some $(x_0, \alpha) \in \mathbb{R}_+^L \times \mathbb{R}$, the sequence of initial conditions $\left\{X^{(\eta)}(0) = \left(x^{(\eta)}(0), z_1^{(\eta)}(0), z_2^{(\eta)}(0)\right)\right\}$ satisfies*

$$\limsup_{\eta\to\infty} \left\| \left(x^{(\eta)}(0), z_1^{(\eta)}(0) - z_2^{(\eta)}(0)\right) - (x_0, \alpha) \right\| \le \epsilon_0.$$

*Define $\sigma_\eta(\epsilon)$ and $\mu_\eta(\epsilon)$ as*

$$\sigma_\eta(\epsilon) \triangleq \inf\left\{t \ge 0 : \left|z_1^{(\eta)}(t) - z_2^{(\eta)}(t)\right| < \epsilon\right\} \quad \text{and} \quad \mu_\eta(\epsilon) \triangleq \inf\left\{t \ge \sigma_\eta(\epsilon) : \left|z_1^{(\eta)}(t) - z_2^{(\eta)}(t)\right| > 2\epsilon\right\}.$$

*Letting $\big(x(t), z(t)\big)$ be the solution of system $\mathcal{S}$, we have the following*

*(A) There exists a constant $C_1 > 0$ such that*

$$\limsup_{\eta\to\infty} \left[ e^{-C_1(\mu_\eta(\epsilon)\wedge T)} \sup_{t\in[0,\mu_\eta(\epsilon)\wedge T]} \left\| \left(x^{(\eta)}(t), z_1^{(\eta)}(t) - z_2^{(\eta)}(t)\right) - \big(x(t), z(t)\big) \right\| \le \epsilon_0 \right].$$

*(B) There exists a constant $C_2 > 0$ such that*

$$\limsup_{\eta\to\infty} \frac{1}{\mu_\eta(\epsilon)\wedge T} \int_0^{\mu_\eta(\epsilon)\wedge T} \left\| \left(x^{(\eta)}(t), z_1^{(\eta)}(t), z_2^{(\eta)}(t)\right) - \big(x(t), z^+(t), z^-(t)\big) \right\| dt \le C_2\epsilon_0.$$

*Moreover the constants $C_1$ and $C_2$ can be chosen to be independent of $\epsilon_0, \epsilon$ and $(x_0, \alpha)$.*

**Proof.** Without loss of generality we can assume that there exists a $\widetilde{x}_0 \in \mathbb{R}_+^L, \widetilde{\theta} > 0$ and $\widetilde{\alpha} \in \mathbb{R}$ such that $X^{(\eta)}(0) = (\widetilde{x}_0, \widetilde{\theta} + \widetilde{\alpha}, \widetilde{\theta})$ for each $\eta$ and

$$\|(\widetilde{x}_0, \widetilde{\alpha}) - (x_0, \alpha)\| \le \epsilon_0.$$

Let $\left(\widetilde{x}(t), \widetilde{z}(t)\right)$ be the solution of system $\mathcal{S}$ but with initial condition $(\widetilde{x}_0, \widetilde{\alpha})$. Due to the Lipchitz nature of functions $F$, $W_1$ and $W_2$ there exists a constant $M > 0$ such that for any $t \ge 0$

$$\left\|\big(\widetilde{x}(t), \widetilde{z}(t)\big) - \big(x(t), z(t)\big)\right\| \le \epsilon_0 e^{Mt}. \tag{17}$$

For convenience we assume that $\mu_\eta(\epsilon) \le T$, but the proof works in the case $\mu_\eta(\epsilon) \ge T$ as well. Observe that

$$\begin{aligned}
&\sup_{t\in[0,\mu_\eta(\epsilon)]} \left\| \left(x^{(\eta)}(t), z_1^{(\eta)}(t) - z_2^{(\eta)}(t)\right) - \big(x(t), z(t)\big) \right\| \\
&\quad\le \sup_{t\in[0,\mu_\eta(\epsilon)]} \left\| \left(x^{(\eta)}(t), z_1^{(\eta)}(t) - z_2^{(\eta)}(t)\right) - \big(\widetilde{x}(t), \widetilde{z}(t)\big) \right\| + \sup_{t\in[0,\mu_\eta(\epsilon)]} \left\| \big(\widetilde{x}(t), \widetilde{z}(t)\big) - \Big(x\big(\mu_\eta(\epsilon)\big), z\big(\mu_\eta(\epsilon)\big)\Big) \right\| \\
&\quad\le \sup_{t\in[0,\mu_\eta(\epsilon)]} \left\| \left(x^{(\eta)}(t), z_1^{(\eta)}(t) - z_2^{(\eta)}(t)\right) - \big(\widetilde{x}(t), \widetilde{z}(t)\big) \right\| + \epsilon_0 e^{M\mu_\eta(\epsilon)}, \qquad (18)
\end{aligned}$$

where the last inequality holds due to (17). From Proposition S2 we know that

$$\limsup_{\eta\to\infty} \sup_{t\in[0,\mu_\eta(\epsilon)]} \left\| \left(x^{(\eta)}(t), z_1^{(\eta)}(t) - z_2^{(\eta)}(t)\right) - \left(\widetilde{x}(t), \widetilde{z}(t)\right) \right\| = 0,$$

and $\sigma_\eta(\epsilon) \to \sigma(\epsilon) \triangleq \inf\{t \geq 0 : |\widetilde{y}(t)| < \epsilon\}$. Without losing generality we can assume that $\sigma_\eta(\epsilon) = \sigma(\epsilon)$ and $\left(x^{(\eta)}(t), z_1^{(\eta)}(t) - z_2^{(\eta)}(t)\right) = \left(\widetilde{x}(t), \widetilde{z}(t)\right)$ for each $\eta$ and $t \in [0, \sigma(\epsilon)]$. Let $\tau_\eta(\epsilon) \triangleq \mu_\eta(\epsilon) - \sigma(\epsilon)$, and for any $t \in [0, \tau_\eta(\epsilon)]$ define

$$\left(\bar{x}^{(\eta)}(t), \bar{z}_1^{(\eta)}(t), \bar{z}_2^{(\eta)}(t)\right) \triangleq \left(\widetilde{x}^{(\eta)}(t+\sigma(\epsilon)), \widetilde{z}_1^{(\eta)}(t+\sigma(\epsilon)), \widetilde{z}_2^{(\eta)}(t+\sigma(\epsilon))\right)$$
$$\left(\bar{x}(t), \bar{z}(t)\right) \triangleq \left(\widetilde{x}(t+\sigma(\epsilon)), \widetilde{z}(t+\sigma(\epsilon))\right).$$

One can see that

$$\sup_{t\in[0,\mu_\eta(\epsilon)]} \left\| \left(x^{(\eta)}(t), z_1^{(\eta)}(t) - z_2^{(\eta)}(t)\right) - \left(\widetilde{x}(t), \widetilde{z}(t)\right) \right\| = \sup_{t\in[0,\tau_\eta(\epsilon)]} \left\| \left(\bar{x}^{(\eta)}(t), \bar{z}_1^{(\eta)}(t) - \bar{z}_2^{(\eta)}(t)\right) - \left(\bar{x}(t), \bar{z}(t)\right) \right\|.$$

Also note that $\lim_{\eta\to\infty} \left(\bar{x}^{(\eta)}(0), \bar{z}_1^{(\eta)}(0), \bar{z}_2^{(\eta)}(0)\right) = \left(\widetilde{x}(\sigma(\epsilon)), \epsilon, 0\right)$ or $(\widetilde{x}\left(\sigma(\epsilon)\right), 0, \epsilon\big)$. From Proposition S1, relation (6) we see that there exists a constant $C > 0$ such that for large $\eta$ we have

$$\sup_{t\in[0,\tau_\eta(\epsilon)]} \left\| \left(\bar{x}^{(\eta)}(t), \bar{z}_1^{(\eta)}(t) - \bar{z}_2^{(\eta)}(t)\right) - \left(\bar{x}(t), \bar{z}(t)\right) \right\| \leq C\epsilon\tau_\eta(\epsilon)e^{C\tau_\eta(\epsilon)}.$$

Using (18) we obtain the following for large $\eta$:

$$\sup_{t\in[0,\mu_\eta(\epsilon)]} \left\| \left(x^{(\eta)}(t), z_1^{(\eta)}(t) - z_2^{(\eta)}(t)\right) - \left(x(t), z(t)\right) \right\| \leq C\epsilon\tau_\eta(\epsilon)e^{C\tau_\eta(\epsilon)} + \epsilon_0 e^{M\mu_\eta(\epsilon)}.$$

Let $\bar{M} \triangleq \max\{C, M\}$. Since $\epsilon \leq \epsilon_0, \tau_\eta(\epsilon) \leq \mu_\eta(\epsilon)$ and $1 + x \leq e^x$ for any $x \geq 0$, we get

$$\sup_{t\in[0,\mu_\eta(\epsilon)]} \left\| \left(x^{(\eta)}(t), z_1^{(\eta)}(t) - z_2^{(\eta)}(t)\right) - \left(x(t), z(t)\right) \right\| \leq \epsilon_0 e^{\bar{M}\mu_\eta(\epsilon)}\left(1 + \bar{M}\mu_\eta(\epsilon)\right) \leq \epsilon_0 e^{2\bar{M}\mu_\eta(\epsilon)},$$

which proves part (A) of the proposition. The proof of part (B) is similar except that supremum must be replaced by time-integral, and instead of relation (6) we use relation (7) in Proposition S1. □

We are now ready to prove Theorem 1. Let $X^{(\eta)}(t) = \left(x^{(\eta)}(t), z_1^{(\eta)}(t), z_2^{(\eta)}(t)\right)$ and $\left(x(t), z(t)\right)$ be as in the statement of this theorem. Without loss of generality we can assume that $X^{(\eta)}(0) = (x_0, a, b)$ for each $\eta$. We first consider the case $a \neq b$. Pick an $\epsilon > 0$. Let $\mu_\eta^{(0)}(\epsilon) = 0$ and for each $i = 1, 2, \ldots$ define $\sigma_\eta^{(i)}(\epsilon)$ and $\mu_\eta^{(i)}(\epsilon)$ as

$$\sigma_\eta^{(i)}(\epsilon) \triangleq \inf\left\{t \geq \mu_\eta^{(i-1)}(\epsilon) : \left|z_1^{(\eta)}(t) - z_2^{(\eta)}(t)\right| < \epsilon\right\} \text{ and } \mu_\eta^{(i)}(\epsilon) \triangleq \inf\left\{t \geq \sigma_\eta^{(i)}(\epsilon) : \left|z_1^{(\eta)}(t) - z_2^{(\eta)}(t)\right| > 2\epsilon\right\}.$$

Let constants $C_1$ and $C_2$ be as in Lemma S3. Let $\epsilon_0 = \epsilon$ and for $i = 1, 2, \ldots$ define

$$\epsilon_i \triangleq \epsilon_{i-1} e^{C_1\left(\mu_\eta^{(i)}(\epsilon)\wedge T - \mu_\eta^{(i-1)}(\epsilon)\wedge T\right)} = \epsilon e^{\sum_{j=1}^{i} C_1\left(\mu_\eta^{(j)}(\epsilon)\wedge T - \mu_\eta^{(j-1)}(\epsilon)\wedge T\right)} = \epsilon e^{C_1\left(\mu_\eta^{(i)}(\epsilon)\wedge T\right)}.$$

Using Proposition S1 for $i = 1$ and Lemma S3 part (A) for $i \geq 2$ we get that for each $i$ we have

$$\sup_{t\in\left[0,\mu_\eta^{(i)}(\epsilon)\wedge T\right]} \left\| \left(x^{(\eta)}(t), z_1^{(\eta)}(t) - z_2^{(\eta)}(t)\right) - \left(x(t), z(t)\right) \right\| \leq \epsilon_i,$$

for large $\eta$. Note that $\epsilon_i < \epsilon e^{C_1 T}$. Letting $\eta \to \infty$ and then $i \to \infty$, and using Monotone Convergence Theorem implies that

$$\limsup_{\eta\to\infty} \sup_{t\in[0,T]} \left\| \left( x^{(\eta)}(t), z_1^{(\eta)}(t) - z_2^{(\eta)}(t) \right) - \left( x(t), z(t) \right) \right\| \leq \epsilon e^{C_1 T},$$

and letting $\epsilon \to 0$ proves the first convergence in Theorem 1.

From part (B) of Lemma S3 we know that for each $i = 1, 2, \ldots$ we have

$$\int_{\mu_\eta^{(i-1)}(\epsilon)\wedge T}^{\mu_\eta^{(i)}(\epsilon)\wedge T} \left\| \left( x^{(\eta)}(t), z_1^{(\eta)}(t), z_2^{(\eta)}(t) \right) - \left( x(t), z^+(t), z^-(t) \right) \right\| dt \leq C_2 \epsilon_{i-1} \left( \mu_\eta^{(i)}(\epsilon) \wedge T - \mu_\eta^{(i-1)}(\epsilon) \wedge T \right),$$

for large $\eta$. This shows that

$$\begin{aligned}
&\int_0^T \left\| \left( x^{(\eta)}(t), z_1^{(\eta)}(t), z_2^{(\eta)}(t) \right) - \left( x(t), z^+(t), z^-(t) \right) \right\| dt \\
&\qquad = \sum_{i=1}^{\infty} \int_{\mu_\eta^{(i-1)}(\epsilon)\wedge T}^{\mu_\eta^{(i)}(\epsilon)\wedge T} \left\| \left( x^{(\eta)}(t), z_1^{(\eta)}(t), z_2^{(\eta)}(t) \right) - \left( x(t), z^+(t), z^-(t) \right) \right\| dt \\
&\qquad \leq \sum_{i=1}^{\infty} C_2 \epsilon_{i-1} \left( \mu_\eta^{(i)}(\epsilon) \wedge T - \mu_\eta^{(i-1)}(\epsilon) \wedge T \right) \\
&\qquad \leq \epsilon C_2 e^{C_1 T} \sum_{i=1}^{\infty} \left( \mu_\eta^{(i)}(\epsilon) \wedge T - \mu_\eta^{(i-1)}(\epsilon) \wedge T \right) \leq \epsilon C_2 e^{C_1 T} T.
\end{aligned}$$

Letting $\eta \to \infty$ and then $\epsilon \to 0$ proves the second convergence in Theorem 1. This completes the proof.

#### S3 Proof of Theorem 2

Let $\bar{X}^{(\eta)} \triangleq \left( \bar{x}^{(\eta)}, \bar{z}_1^{(\eta)}, \bar{z}_2^{(\eta)} \right)$ denote a non-negative fixed point of system $\mathcal{S}_\eta$, with $\eta > 0$. Hence, we have

$$\begin{cases} F\left(\bar{X}^{(\eta)}\right) = 0 \\ \dfrac{1}{\eta} W_1\left(\bar{X}^{(\eta)}\right) - \bar{z}_1^{(\eta)} \bar{z}_2^{(\eta)} = 0 \\ \dfrac{1}{\eta} W_2\left(\bar{X}^{(\eta)}\right) - \bar{z}_1^{(\eta)} \bar{z}_2^{(\eta)} = 0 \end{cases} \iff \begin{cases} F\left(\bar{X}^{(\eta)}\right) = 0 \\ W_1\left(\bar{X}^{(\eta)}\right) - W_2\left(\bar{X}^{(\eta)}\right) = 0 \\ \dfrac{1}{\eta} W_2\left(\bar{X}^{(\eta)}\right) - \bar{z}_1^{(\eta)} \bar{z}_2^{(\eta)} = 0. \end{cases} \tag{19}$$

Since the limit of $\bar{X}^{(\eta)}$ as $\eta \to \infty$ is not always assumed to exist, then we only consider convergent sub-sequences with limiting points. More precisely, for each $i = 1, 2, \cdots$, let $\{\eta_k^i\}_{k\in\mathbb{N}}$ be a sequence with $\eta_k^i > 0$ and $\lim_{k\to\infty} \eta_k^i = \infty$ such that $\bar{X}^i = (\bar{x}^i, \bar{z}_1^i, \bar{z}_2^i) \triangleq \lim_{k\to\infty} \bar{X}^{(\eta_k^i)}$ is a limiting fixed point along the convergent sub-sequence $\left\{\bar{X}^{(\eta_k^i)}\right\}_{k\in\mathbb{N}}$. Taking the limit of (19) along said sub-sequence yields

$$\begin{cases} \lim_{k\to\infty} F\left(\bar{X}^{(\eta_k^i)}\right) = 0 \\ \lim_{k\to\infty} W_1\left(\bar{X}^{(\eta_k^i)}\right) - \lim_{k\to\infty} W_2\left(\bar{X}^{(\eta_k^i)}\right) = 0 \\ \lim_{k\to\infty} \dfrac{1}{\eta_k^i} W_2\left(\bar{X}^{(\eta_k^i)}\right) - \lim_{k\to\infty} \bar{z}_1^{(\eta_k^i)} \bar{z}_2^{(\eta_k^i)} = 0 \end{cases} \implies \begin{cases} F\left(\bar{X}^i\right) = 0 \\ W_1\left(\bar{X}^i\right) - W_2\left(\bar{X}^i\right) = 0 \\ \bar{z}_1^i \bar{z}_2^i = 0, \end{cases} \tag{20}$$

which follows from the Lipschitz assumption on $F, W_1$ and $W_2$ that allows us to move the limits inside the functions and exploit the fact that $W_j\left(\bar{X}^{(\eta_k^i)}\right)$ for $j = 1, 2$ remain finite in the limit as $k \to \infty$. Now define $\bar{z}^i \triangleq \bar{z}_1^i - \bar{z}_2^i$. Then, invoking the non-negativity assumption of the fixed points, we have

$$\begin{cases} \bar{z}_1^i \bar{z}_2^i = 0 \\ \bar{z}_1^i, \bar{z}_2^i \geq 0 \end{cases} \implies \begin{cases} \bar{z}_1^i = \max(\bar{z}^i, 0) = \left(\bar{z}^i\right)^+ \\ \bar{z}_2^i = \max(-\bar{z}^i, 0) = \left(\bar{z}^i\right)^- . \end{cases} \tag{21}$$

Substituting in (20) yields

$$\begin{cases} F\left(\bar{x}^i, \left(\bar{z}^i\right)^+, \left(\bar{z}^i\right)^-\right) = 0 \\ W_1\left(\bar{x}^i, \left(\bar{z}^i\right)^+, \left(\bar{z}^i\right)^-\right) - W_2\left(\bar{x}^i, \left(\bar{z}^i\right)^+, \left(\bar{z}^i\right)^-\right) = 0, \end{cases} \tag{22}$$

which are exactly the equations of the fixed points for system $\mathcal{S}$ and thus completing the proof.

#### S4 Proof of Corollary 1

Theorem 2 establishes that all convergent sub-sequences, as $\eta \to \infty$, of non-negative fixed points of system $\mathcal{S}_\eta$ yield a subset of the fixed points of system $\mathcal{S}$. Hence, if there is only one fixed point for system $\mathcal{S}$, then there exists only one such convergent sub-sequence. The second part of the corollary is the contrapositive of Theorem 2.

#### S5 Proof of Lemma 2

The closed-loop dynamics describing the block diagram of Fig. 5(b) are given by the following set of differential equations.

$$\begin{array}{rll} \textbf{Controlled Process} & y = \mathcal{P}_\Delta(u) \quad \text{e.g.} & \begin{cases} \dot{x} = f_\Delta(x,u) \\ y = g_\Delta(x,u) \end{cases} \\ \textbf{Error} & e = r_{\text{in}} - w & \\ \textbf{Integrator} & \dot{v} = K_I e & \\ \textbf{Actuator} & u = \psi_a(v) & \\ \textbf{Sensor} & w = \psi_s(y). & \end{array} \tag{23}$$

Thus the dynamics and the steady-state values, if they exist, satisfy

$$\begin{cases} y = \mathcal{P}_\Delta\big(\psi_a(v)\big) \\ \dot{v} = K_I\big(r_{\text{in}} - \psi_s(y)\big) \end{cases} \implies \begin{cases} \bar{\mathcal{P}}_{\bar{\Delta}}\big(\psi_a(\bar{v})\big) = \bar{y} \\ \psi_s(\bar{y}) = r_{\text{in}}. \end{cases}$$

*Sufficiency.* Let $r_{\text{in}} \in \text{range}(\psi_s)$, then there exists a $\bar{y} = r_{\text{out}}$ such that $\psi_s(r_{\text{out}}) = r_{\text{in}}$, and in fact it is unique since $\psi_s$ is strictly monotonically increasing. To this end, we can write $\bar{y} = r_{\text{out}} = \psi_s^{-1}(r_{\text{in}})$. Let $r_{\text{out}} \in \mathcal{R}_{\text{out}}$, then the supporting input exists and is given by $\bar{u} = \bar{\mathcal{P}}_{\bar{\Delta}}^{-1}(r_{\text{out}}) = \bar{\mathcal{P}}_{\bar{\Delta}}^{-1} \circ \psi_s^{-1}(r_{\text{in}})$. Furthermore, if the actuator does not saturate, then $\bar{u} \in \text{range}(\psi_a)$ which implies that there exists a $\bar{v}$ such that $\psi_a(\bar{v}) = \bar{u}$. It is in fact unique since $\psi_a$ is strictly monotonically increasing, and finally we can write $\bar{v} = \psi_a^{-1} \circ \bar{\mathcal{P}}_{\bar{\Delta}}^{-1} \circ \psi_s^{-1}(r_{\text{in}})$.

*Necessity.* This can be straightforwardly established using the contra-positive, that is, if any of the conditions are not satisfied, then the fixed point does not exist with a feasible supporting input. In fact, if $r_{\text{out}} \notin \mathcal{R}_{\text{out}}$ then $\bar{u} \notin \mathbb{U}$. Otherwise, if $\bar{\mathcal{P}}_{\bar{\Delta}}^{-1}(r_{\text{out}}) \notin \text{range}(\psi_a)$ and/or $r_{\text{in}} \notin \text{range}(\psi_s)$, then $\bar{v}$ and/or $\bar{y}$ do not exist, respectively.

#### S6 Model Reduction in the Stochastic Setting

We now consider the same problem in the case where the dynamics are given by a continuous-time Markov chain over the non-negative integer lattice. Suppose we have a well-stirred system consisting of $L$ species. The state at any time can be described by a vector $x \in \mathbb{N}_0^L$. There are $K$ reaction channels, and when the state is $x$ the $k$-th reaction channel fires at rate $\lambda_k(x)$ and displaces the state to $(x + \zeta_k)$ for some $\zeta_k \in \mathbb{Z}^L$. We assume that for any $k = 1, \ldots, K$ and state $x \in \mathbb{N}_+^L$, if $\lambda_k(x) > 0$ then $(x + \zeta_k) \in \mathbb{N}_0^L$. This ensures that the trajectories of the CTMC lie in the non-negative integer orthant $\mathbb{N}_0^L$. We now augment the system with two more species $\mathbf{Z}_1$ and $\mathbf{Z}_2$ and the following five additional reactions:

| Reaction | Propensity | Stoichiometry Vector |
|---|---|---|
| $\varnothing \longrightarrow \mathbf{Z}_1$ | $W_{1+}(x,z_1,z_2)$ | $(\mathbf{0},1,0)$ |
| $\mathbf{Z}_1 \longrightarrow \varnothing$ | $W_{1-}(x,z_1,z_2)$ | $(\mathbf{0},-1,0)$ |
| $\varnothing \longrightarrow \mathbf{Z}_2$ | $W_{2+}(x,z_1,z_2)$ | $(\mathbf{0},0,1)$ |
| $\mathbf{Z}_2 \longrightarrow \varnothing$ | $W_{2-}(x,z_1,z_2)$ | $(\mathbf{0},0,-1)$ |
| $\mathbf{Z}_1+\mathbf{Z}_2 \longrightarrow \varnothing$ | $\eta z_1 z_2$ | $(\mathbf{0},-1,-1)$ |

Observe that the functions $W_j$, for $j=1,2$ from (1) are now decomposed into two non-negative components $W \triangleq W_{j+} - W_{j-}$ such that the dynamics remain in the non-negative orthant. We also suppose that all the propensity functions $\lambda_k$, for $k=1,\cdots,K$ can depend on the copy-numbers of these two species $\mathbf{Z}_1$ and $\mathbf{Z}_2$. The random time-change representation for the overall dynamics is given by

$$X^{(\eta)}(t) = X^{(\eta)}(0) + \sum_{k=1}^{K} Y_k\left(\int_0^t \lambda_k\left(X^{(\eta)}(s), Z_1^{(\eta)}(s), W_2^{(\eta)}(s)\right) ds\right)\zeta_k$$

$$Z_1^{(\eta)}(t) = Z_1^{(\eta)}(0) + \sum_{\ell\in\{+,-\}} \ell Y_{K+1}^{\ell}\left(\int_0^t W_{1l}\left(X^{(\eta)}(s), Z_1^{(\eta)}(s), Z_2^{(\eta)}(s)\right) ds\right) - Y_{K+3}\left(\eta\int_0^t Z_1^{(\eta)}(s)Z_2^{(\eta)}(s)ds\right)$$

$$Z_2^{(\eta)}(t) = Z_2^{(\eta)}(0) + \sum_{\ell\in\{+,-\}} \ell Y_{K+2}^{\ell}\left(\int_0^t W_{2\ell}\left(X^{(\eta)}(s), Z_1^{(\eta)}(s), Z_2^{(\eta)}(s)\right) ds\right) - Y_{K+3}\left(\eta\int_0^t Z_1^{(\eta)}(s)Z_2^{(\eta)}(s)ds\right),$$

where $Y_1,\cdots,Y_K, Y_{K+1}^{\pm}, Y_{K+2}^{\pm}$ and $Y_{K+3}$ are $K+5$ independent unit-rate Poisson processes.

For large values of $\eta$, the reaction $\mathbf{Z}_1+\mathbf{Z}_2 \longrightarrow \varnothing$ operates on a *faster* timescale than other reactions. We shall use the results in [2] to analyze the limiting behavior of the overall dynamics $\left(X^{(\eta)}(t), Z_1^{(\eta)}(t), Z_2^{(\eta)}(t)\right)_{t\geq 0}$ as $\eta\to\infty$. Let $Z^{(\eta)}(t) \triangleq Z_1^{(\eta)}(t) - Z_2^{(\eta)}(t)$ and note that the dynamics of the process $\left(X^{(\eta)}(t), Z^{(\eta)}(t)\right)_{t\geq 0}$ is unaffected by the fast reaction. Next, define a random measure on $\mathbb{N}_0^2 \times [0,\infty)$ by

$$V^{(\eta)}(C\times[0,t]) = \int_0^t \mathbb{1}_C\left(Z_1^{(\eta)}(s), Z_2^{(\eta)}(s)\right) ds.$$

For any $z\in\mathbb{Z}$, define an operator that acts on some bounded function $f$ as

$$\mathbb{C}^z f(z_1,z_2) = z_1 z_2\Big(f(z_1-1,z_2-1) - f(z_1,z_2)\Big).$$

This operator is the generator of the dynamics of species $\mathbf{Z}_1$ and $\mathbf{Z}_2$ due to the fast reaction $\mathbf{Z}_1+\mathbf{Z}_2 \longrightarrow \varnothing$, when initial difference in their copy numbers is $z$. Note that the Markovian dynamics generated by this dynamics are ergodic with unique stationary distribution given by

$$\pi_z(z_1,z_2) = \begin{cases} 1 & \text{if } z_1 = z, z_2 = 0 \text{ and } z\geq 0 \\ 1 & \text{if } z_1 = 0, z_2 = -z \text{ and } z<0 \\ 0 & \text{otherwise.}\end{cases}$$

For each $k=1,\ldots,K$ and $j=1,2$ define

$$\widehat{\lambda}_k(x,z) \triangleq \mathbb{E}_{\pi_z}\left[\lambda_k(x,Z_1,Z_2)\right] = \sum_{\mathbb{N}_0^2} \lambda_k(x,z_1,z_2)\pi_z(z_1,z_2) = \lambda_k\left(x,z^+,z^-\right)$$

$$\widehat{W}_{j\pm}(x,z) \triangleq \mathbb{E}_{\pi_z}\left[W_{j\pm}(x,Z_1,Z_2)\right] = \sum_{\mathbb{N}_0^2} W_{j\pm}(x,z_1,z_2)\pi_z(z_1,z_2) = W_{j\pm}\left(x,z^+,z^-\right).$$

As the Markov process corresponding to generator $\mathbb{C}^z$ is ergodic with stationary distribution $\pi_z$, we can expect that if we have $\left(X^{(\eta)}, Z^{(\eta)}, V^{(\eta)}\right) \to (X,Z,V)$ as $\eta\to\infty$, then the limiting occupation measure $V$ has the form

$$V(dy\times ds) = \pi_{Z(s)}(dy)ds. \tag{24}$$

Moreover since for each $\lambda_k$ and $W_{j\pm}$ we have

$$\int_0^t \lambda_k\left(X^{(\eta)}(s), Z_1^{(\eta)}(s), Z_2^{(\eta)}(s)\right) ds = \int_0^t \sum_{\mathbb{N}_0^2} \lambda_k\left(X^{(\eta)}(s), z_1, z_2\right) V^{(\eta)}\left(\{z_1, z_2\} \times ds\right) \underset{\eta\to\infty}{\longrightarrow} \int_0^t \widehat{\lambda}_k\Big(X(s), Z(s)\Big) ds,$$

and

$$\int_0^t W_{j\pm}\left(X^{(\eta)}(s), Z_1^{(\eta)}(s), Z_2^{(\eta)}(s)\right) ds = \int_0^t \sum_{\mathbb{N}_0^2} W_{j\pm}\left(X^{(\eta)}(s), z_1, z_2\right) V^{(\eta)}\left(\{z_1, z_2\} \times ds\right) \underset{\eta\to\infty}{\longrightarrow} \int_0^t \widehat{W}_{j\pm}\Big(X(s), Z(s)\Big) ds,$$

the limiting process $(X(t), Z(t))_{t\geq 0}$ has the random time-change representation given by

$$X(t) = X(0) + \sum_{k=1}^{K} Y_k\left(\int_0^t \widehat{\lambda}_k\Big(X(s), Z(s)\Big) ds\right)\zeta_k \tag{25}$$

$$\text{and} \quad Z(t) = Z(0) + \sum_{\ell\in\{+,-\}} \ell Y_{K+1}^{\ell}\left(\int_0^t \widehat{W}_{1\ell}\Big(X(s), Z(s)\Big) ds\right) - \sum_{\ell\in\{+,-\}} \ell Y_{K+2}^{\ell}\left(\int_0^t \widehat{W}_{2\ell}\Big(X(s), Z(s)\Big) ds\right).$$

We now state the convergence result whose proof follows from Theorem 5.1 in [2].

**Proposition S3** *Let the process* $\left(X^{(\eta)}, Z^{(\eta)}\right)_{t\geq 0}$ *and the occupation measure* $V^{(\eta)}$ *be as defined above. Suppose* $\left(X^{(\eta)}(0), Z^{(\eta)}(0)\right) \longrightarrow \left(X(0), U(0)\right)$ *as* $\eta \to \infty$*. We then have* $\left(X^{(\eta)}, Z^{(\eta)}, V^{(\eta)}\right) \longrightarrow (X, Z, V)$ *where* $V$ *is given by* (24) *and the process* $\left(X(t), Z(t)\right)_{t\geq 0}$ *is given by* (25)*.*

For a function $f : \mathbb{R}_+^L \times \mathbb{R} \to \mathbb{R}$ the convergence $\left(X^{(\eta)}(t), Z^{(\eta)}(t)\right)_{t\geq 0} \underset{\eta\to\infty}{\longrightarrow} \left(X(t), Z(t)\right)_{t\geq 0}$ implies that

$$\lim_{\eta\to\infty} \sup_{t\in[0,T]} \left|\mathbb{E}\left[f\left(X^{(\eta)}(t), Z_1^{(\eta)}(t) - Z_2^{(\eta)}(t)\right)\right] - \mathbb{E}\Big[f\Big(X(t), Z(t)\Big)\Big]\right| = 0.$$

for any $T > 0$. Since we also have convergence of the occupation measure we can also conclude that for any $h : \mathbb{R}_+^{L+2} \to \mathbb{R}$ we have

$$\lim_{\eta\to\infty} \int_0^T \left|\mathbb{E}\left[h\left(X^{(\eta)}(t), Z_1^{(\eta)}(t), Z_2^{(\eta)}(t)\right)\right] - \mathbb{E}\Big[h\Big(X(t), Z^+(t), Z^-(t)\Big)\Big]\right| dt = 0$$

for any $T > 0$.

### S7 Supplementary Figures

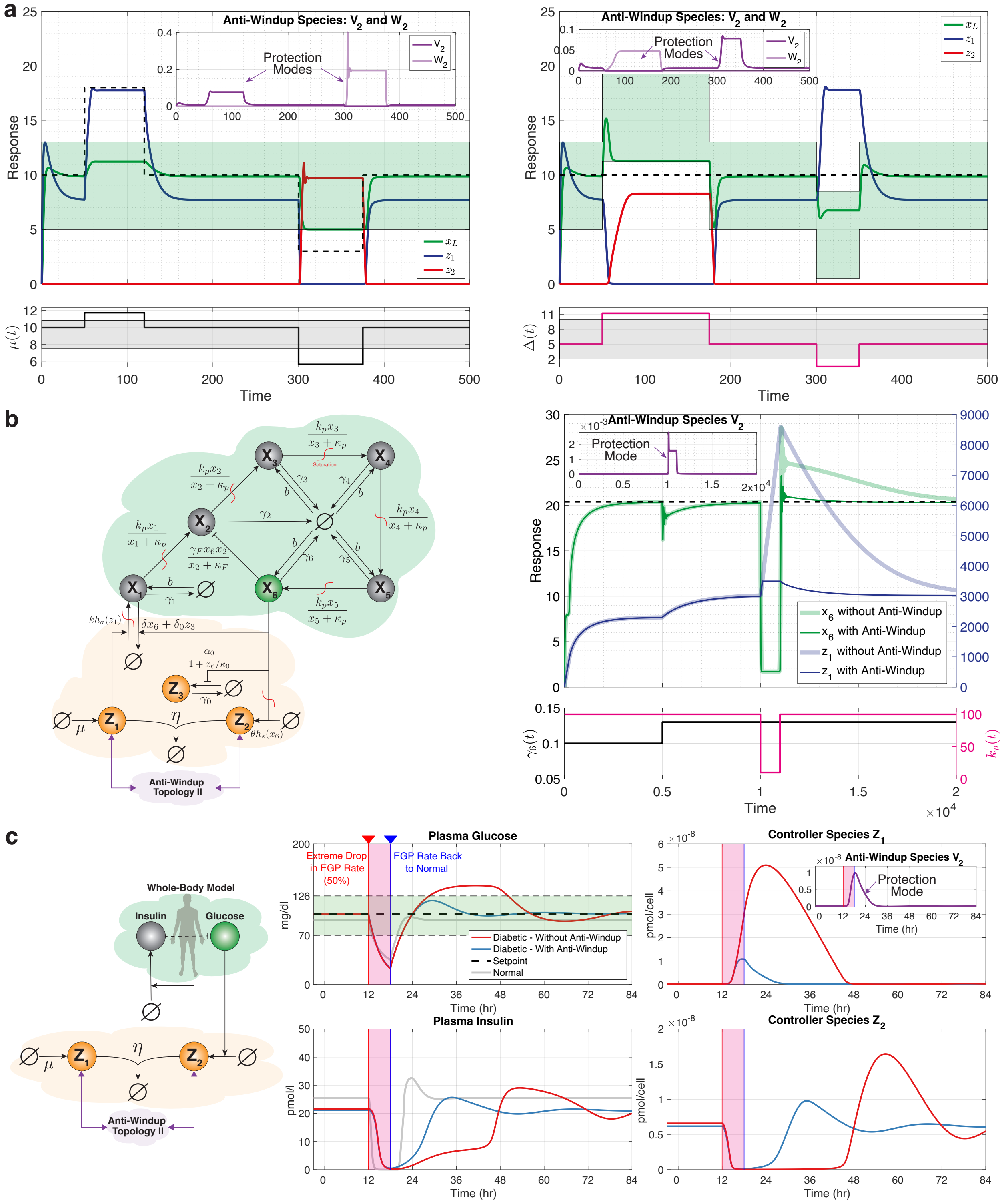


Figure S1: **Reproducing Fig. 10 using anti-windup topology II**. All parameters are the same as those in Fig. 10 except $v_0 = 20, w_0 = 10$ in panel (a), and the functions from Fig. 8 are given by $h_i(v_j) = bv_j$ with $b = 100\mu\text{mol}^{-1}\text{hr}^{-1}$.

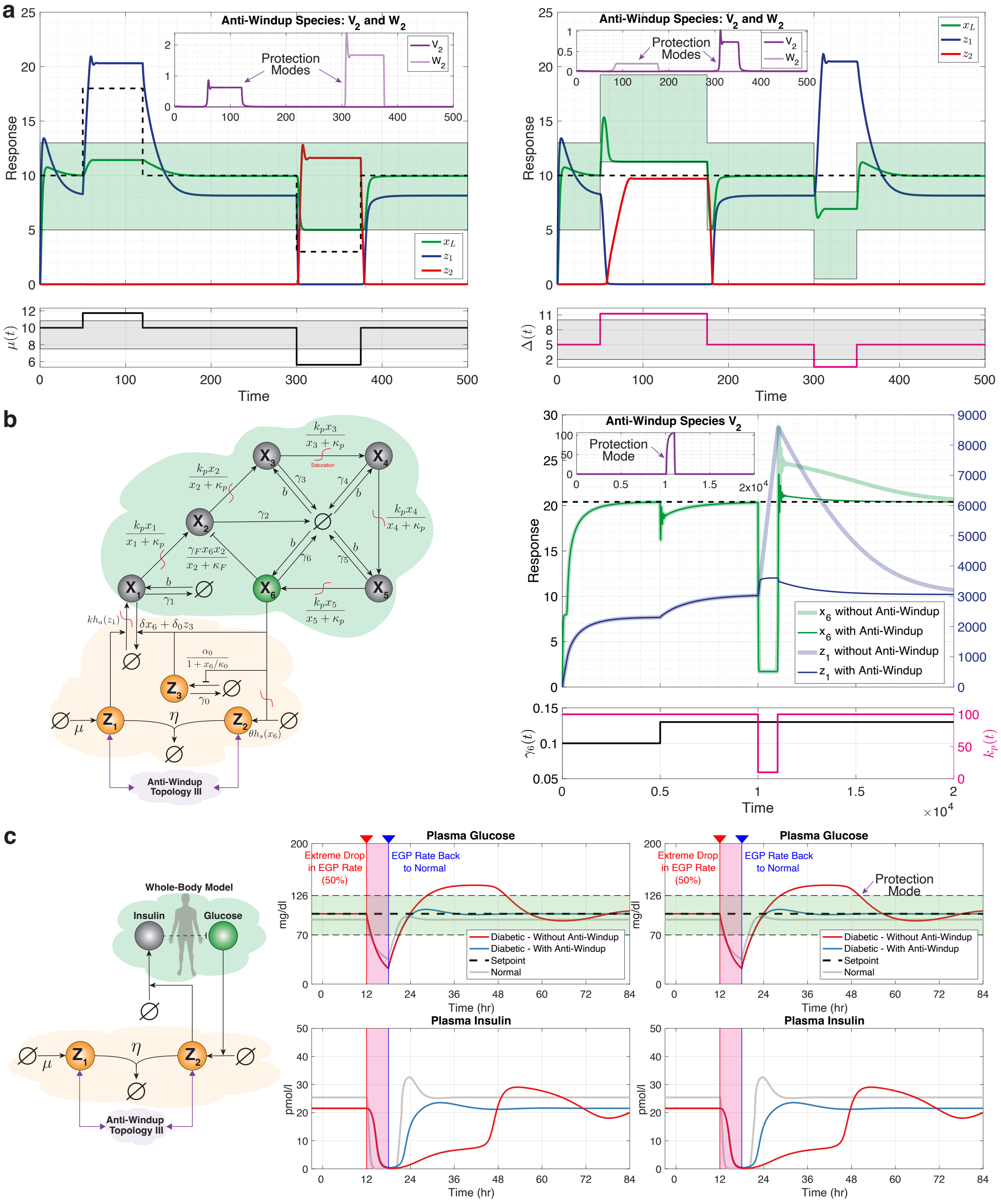


Figure S2: **Reproducing Fig. 10 using anti-windup topology III**. All parameters are the same as those in Fig. 10 except $v_0 = 20, w_0 = 10$ in panel (a). However, the functions from Fig. 8 are now given by $h_i(v_j) = \frac{\alpha}{1+v_j/\kappa}$ with $\alpha = 1, \kappa = 5$ in panel (a), $\alpha = 1, \kappa = 10$ in panel (b) and $\alpha = 1, \kappa = 0.01\mu$mol in panel (c).

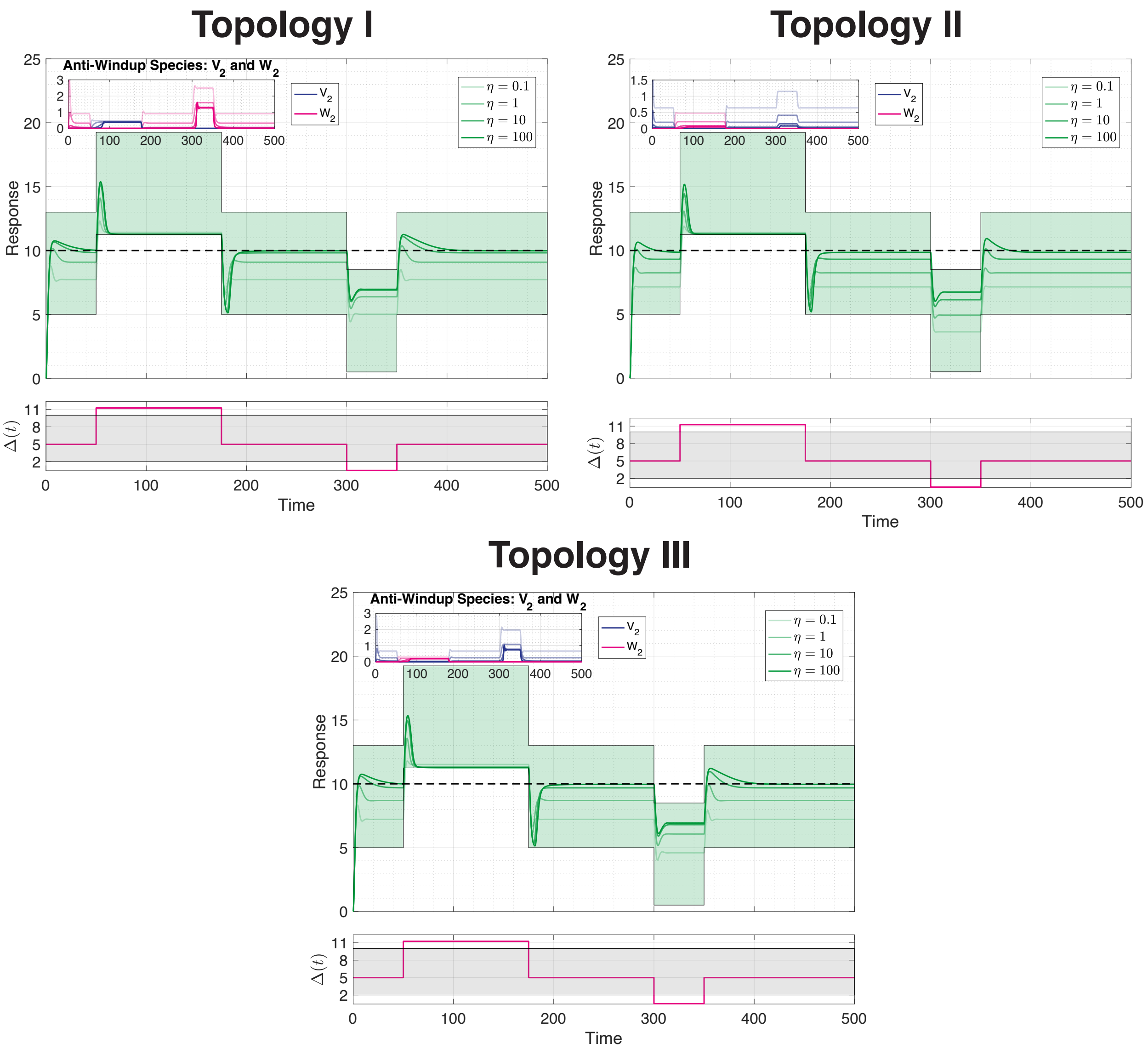


Figure S3: **The effect of the sequestration rates of the three anti-windup topologies**. All parameters are the same as those in Fig. 10(a) for Topology I, Fig. S1(a) for Topology II and Fig.S2(a) for Topology III. The only difference is that $\eta_v = \eta_w$ take the values $\{10^{-1}, 10^0, 10^1, 10^2\}$. For small sequestration rates $\eta_v$ and $\eta_w$, the anti-windup species $\mathbf{V_2}$ and $\mathbf{W_2}$ will have non-zero values even in non-protection modes and thus may interfere with the integrator. As a result, steady-state errors may emerge even when windup is absent. This can be mitigated by having strong sequestration reactions. Observe that for all three topologies, the steady-state error in the non-protection mode decreases to become negligible as $\eta_v$ and $\eta_w$ are increased.

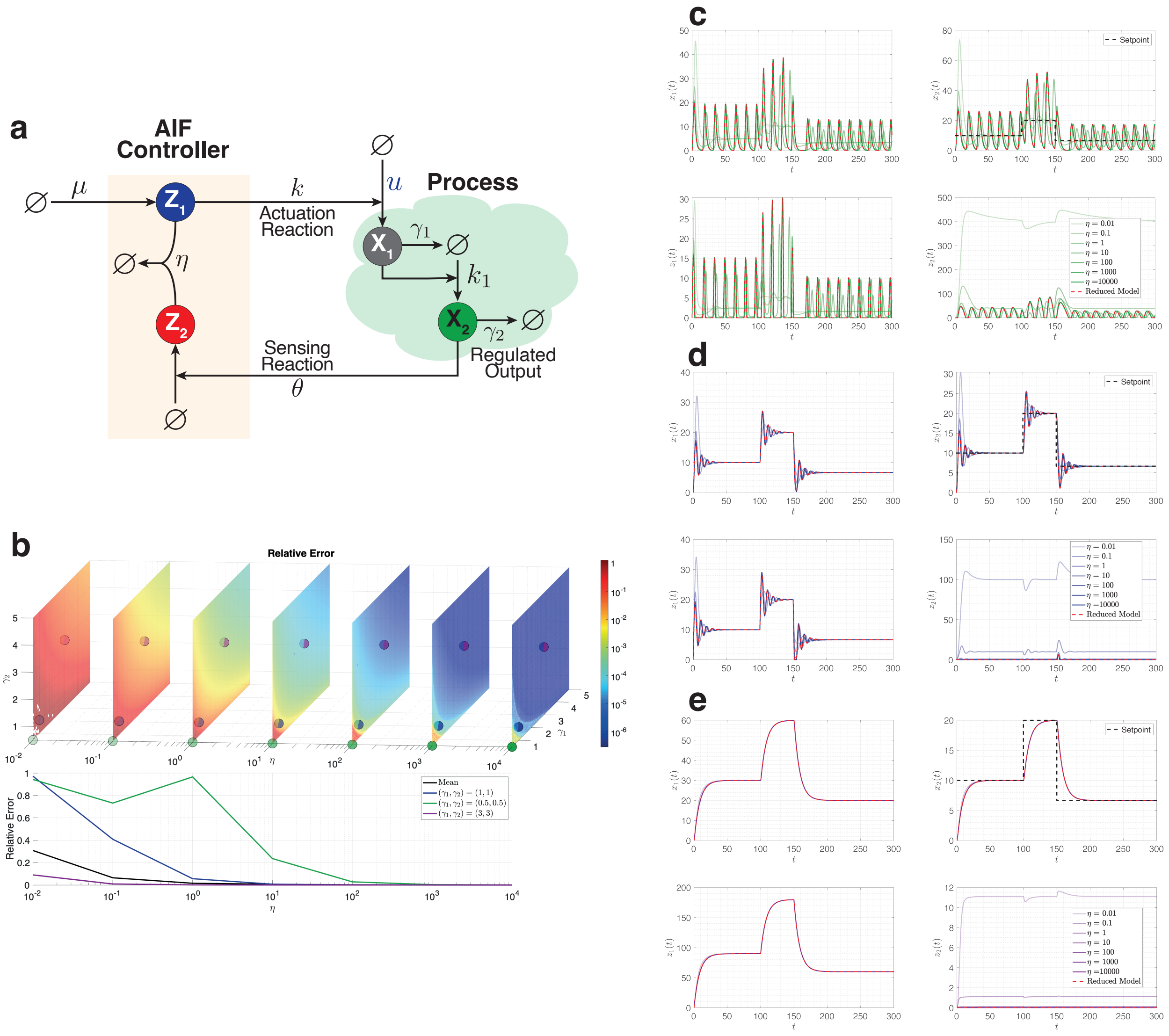


Figure S4: **Deterministic Model Reduction: A Numerical Validation.** (a) The simulations in this figure considers a closed-loop network that integrates the process highlighted in Example 1 with the antithetic integral controller. We set parameters as follows: $\mu = 10, k = \theta = k_1 = 1$. We adjust the values of $\gamma_1$ and $\gamma_2$ between 0.5 and 5, and increase $\eta$ logarithmically, ranging from 0.01 to $10^4$. (b) An examination of the relative error between the full and reduced model, as supported by Theorem 1. These errors, illustrated as intensity plots, span values of $\gamma_1$ and $\gamma_2$ for every $\eta$. Specifically, we calculate the $L^2$-norm over time and states $(x_1, x_2, z_1, z_2)$ of the difference in response between the reduced and full model. This is subsequently normalized by the $L^2$-norm of the full model's response. The graph at the bottom presents the average relative error across $(\gamma_1, \gamma_2)$ values and highlights three specific pairs of $(\gamma_1, \gamma_2)$. Panels (c), (d), and (e) provide detailed response trajectories of the full model for these three pairs of $(\gamma_1, \gamma_2)$. As we elevate $\eta$, these trajectories approach the response of the reduced model, which is depicted in red.

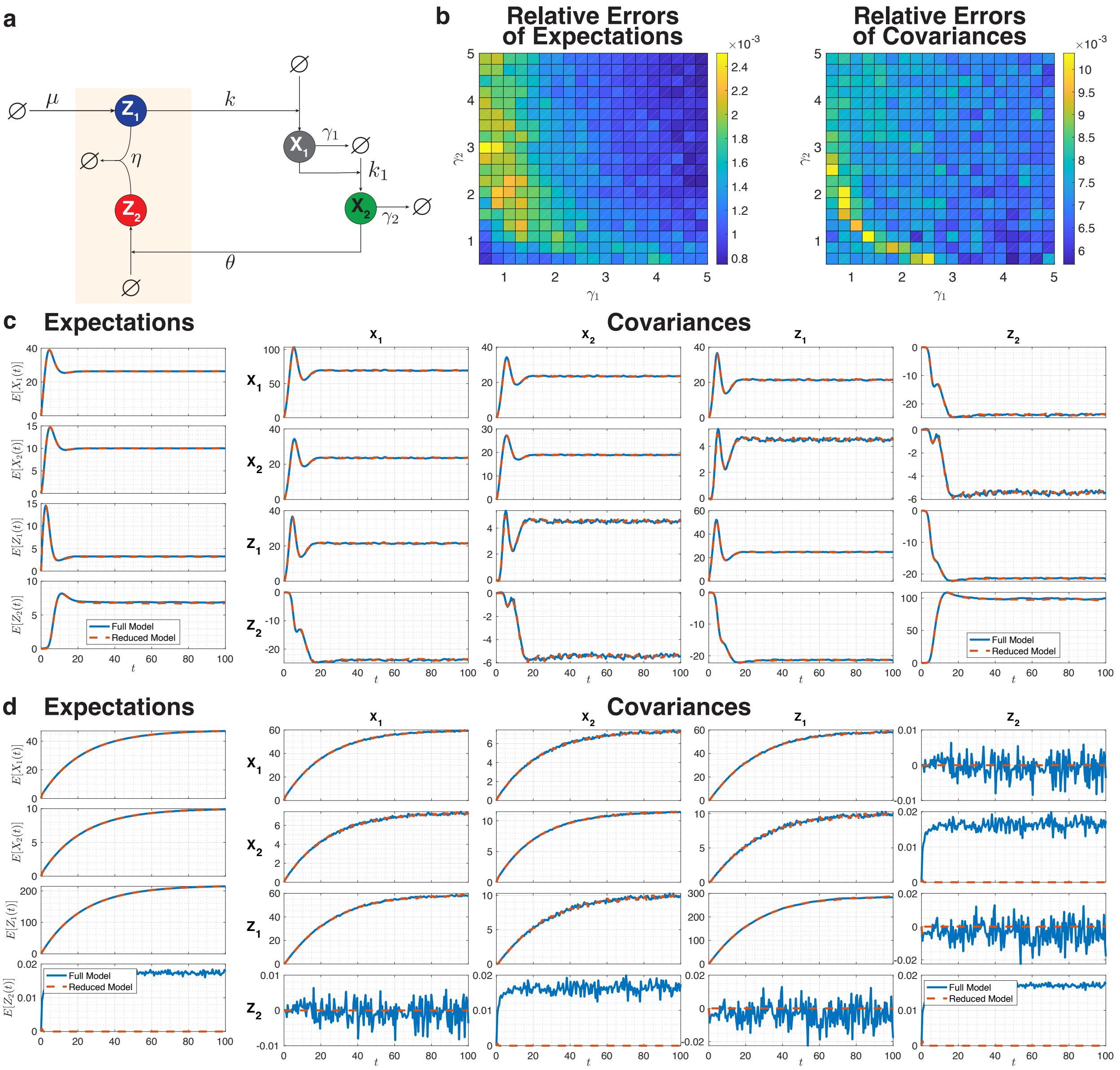


Figure S5: **Stochastic Model Reduction: A Numerical Validation.** (a) The stochastic simulations in this figure consider a closed-loop network that integrates the process highlighted in Example 1 with the antithetic integral controller. We set parameters as follows: $\mu = k_0 = 10, k = \theta = k_1 = 1, \eta = 10^4$. We adjust the values of $\gamma_1$ and $\gamma_2$ between 0.5 and 5. (b) An examination of the relative error between the full and reduced model at the level of the expectations (left) and covariances (right), as supported by Proposition S3. These errors, illustrated as intensity plots, span values of $\gamma_1$ and $\gamma_2$. Specifically, we calculate the $L^2$-norm over time and states $(x_1, x_2, z_1, z_2)$ of the difference in means and covariances between the reduced and full model. This is subsequently normalized by the $L^2$-norm of the full model's mean and covariance. Panels (c) and (d) provide detailed response trajectories of the expectations and covariances of the full and reduced models for two pairs of $(\gamma_1, \gamma_2) = (0.5, 2.5)$ and $(\gamma_1, \gamma_2) = (4.5, 4.5)$. The simulations demonstrate that the reduced model captures the first two moments of the full model. The small remaining errors are due to a finite $\eta = 10^4$ and due to noise induced by a limited number of trajectories ($N = 10^5$) generated by Gillespie's stochastic simulation algorithm to compute the dynamics of the moments.
